# A long-term organotypic slice culture model of patient high-grade glioma preserving tumor heterogeneity and architecture, and specificity of CAR T cell targeting

**DOI:** 10.64898/2026.10.04.756617

**Authors:** Simiao Wang, Ching Ouyang, Heather N. Zook, Massimo D’Apuzzo, Hsun Teresa Ku, Xiwei Wu, Angela S. Baker, Yue Hao, Blake Brewster, Michael Berens, Behnam Badie, Christine E. Brown, Michael E. Barish

## Abstract

**Background:** High grade glioma (HGG), encompassing IDH-wildtype glioblastoma (CNS WHO grade 4), diffuse midline glioma (CNS WHO grade 4), oligodendroglioma (IDH-mutant and 1p/19q-codeleted) CNS WHO grade 3, together with IDH-mutant astrocytoma of CNS grade 3 and grade 4, represents the most lethal group of adult diffuse gliomas. Despite significant efforts, obtaining improvements in patient survival has been limited, in part, by difficulties in comprehensive ex vivo modeling of the complexities of HGG. Although organotypic slice cultures and other platforms have shown promise, existing models often fail to capture the intrinsic cellular heterogeneity, microenvironment, and architecture of HGG, restricting investigations into their fundamental biology as well as design of novel therapeutics.

**Methods:** We developed a patient-derived HGG organotypic slice culture (GOSC) platform demonstrating long-term preservation of intrinsic tumor characteristics. Vibratome-sectioned slices were cultured in defined medium under conditions approximating those in situ: reduced oxygen (4%), low glucose (2.5 mM), and no added growth factors. The characteristics of GOSCs after 28 days in culture, including expression of immunotherapy targets, were compared to parental tumors using histological staining, immunofluorescence, multiplexed PhenoCycler (CODEX) imaging, bulk RNA-seq expression profiling, and CAR T cell killing assays.

**Results:** GOSCs preserved the histopathology, cell heterogeneity, tumor microenvironment, and tissue architecture of their parental tumors throughout at least 28 days in culture. Importantly, GOSCs retained parental patterns of immunotherapy target expression, and CAR T cells co-cultured with GOSCs demonstrated target-specific killing consistent with parental tumors.

**Conclusions:** This ex vivo GOSC platform offers long-term preservation of the cellular and structural properties of parental HGG tumors, thereby providing a robust tool for extended preclinical studies, and facilitating development and evaluation of advanced tumor-targeting strategies.

## INTRODUCTION

High grade glioma (HGG), encompassing IDH-wildtype glioblastoma (CNS WHO grade 4), diffuse midline glioma (CNS WHO grade 4), oligodendroglioma (IDH-mutant and 1p/19q-codeleted) CNS WHO grade 3, together with IDH-mutant astrocytoma of CNS grade 3 and grade 4, represents the most lethal group of adult diffuse gliomas, and are the most common brain cancers in adults [1]. Despite intense efforts, improvements in patient survival have remained modest [2]. A significant barrier to advancing novel therapies has been the paucity of ex vivo models preserving the full complexity of the disease.

HGG is characterized by extensive structural and cellular heterogeneity, and a complex tumor microenvironment, all contributing to therapy resistance and eventual tumor recurrence. HGG tumors contain diverse cell types, including neoplastic cells resembling astrocytes, mesenchymal cells, oligodendrocyte progenitor cells, neuronal progenitors, glioblastoma stem-like cells [3, 4], along with abnormal neovascularization [5–7]. Co-existing within these tumors are normal neurons, astrocytes, and oligodendrocytes, along with radial glia neural progenitors, immune cells, mural cells, and endothelial cells [3, 8–11]. Of particular significance, the non-neoplastic cells in the perivascular niche facilitate crosstalk between vasculature and tumor cells, sustaining the survival and self-renewal of glioma stem-like cells [12, 13]. Together, these components play critical roles in tumor development and progression [14–23] and impact therapeutic efficacies [24].

While efforts to develop 3-dimensional organoid models of human HGG have had some success in advancing basic research and therapeutic strategies (reviewed in [25] [26] [27] [28]; see **Supplemental Table S2**), in many instances tissue dissociation or microdissection has limited preservation of the spatial arrangements and overall complexity of HGG tumors [29]. Additionally, non-physiological growth conditions have not always favored survival of vasculature and immune cells contributing to the tumor microenvironment.

Some efforts to better preserve tissue architecture and cell composition of HGG tumors have employed organotypic slice cultures [30] that when applied to patient HGG tissues [31–33] (**Supplemental Table S2)** maintain some structural and cellular heterogeneity. However, short lifetimes and limited fidelity of these cultures have hampered their utility in developing and evaluating emerging therapeutic strategies.

We hypothesized that organotypic culture of patient HGG slices grown under physiologically relevant conditions (4-5% O_2_ [34, 35] and 2.5-5 mM glucose [36]) would better preserve the characteristics of parental tumors. Here, we report on a human HG<u>G O</u>rganotypic <u>S</u>lice <u>C</u>ulture (GOSC) model offering long-term maintenance of tumor architecture, heterogeneity, and microenvironment, as multiple approaches confirm that for up to 28 days in culture GOSCs retain the structural and cellular complexity of their parental tumors. Additionally, we validate immunotherapy target-specific killing and demonstrate translational potential of GOSCs using previously established IL13Rα2– and chlorotoxin (CLTX)-directed CAR T cells [37–39] — both currently in clinical trials (NCT02208362, NCT04214392) [40, 41]. While some previous ex vivo HGG models have preserved important aspects of HGG structure and composition, these are retained together in our GOSC model. A comparison of the GOSC model with models including GLICO, GBO, IBTO, and others is presented in **Supplemental Table S2**. Results described here establish our long-term GOSC model as a valuable platform for studying the fundamental biology of HGG, and for developing and evaluating over relatively long term the complex therapies that will be required to advance treatment for patients with HGG.

## METHODS

### Generation of the HGG organotypic tumor slice cultures

Samples of freshly resected newly diagnosed or recurrent HGG (**Supplemental Table S1**) were collected in chilled H+GPSA medium [42], composed of Hibernate A (TransnetYX), 1x GlutaMax (Gibco), 1x Penicillin-Streptomycin (Gibco), and 1x Amphotericin B (Gibco). Before any processing, samples were delivered to the City of Hope Surgical Pathology Core for documentation. The use of de-identified patient tissues was approved by the City of Hope Institutional Review Board (IRB #07074).

Tumor pieces (∼0.5 to 1.0 cm diameter) were kept on ice in H+GPSA medium and transferred to the laboratory for vibratome sectioning (Leica VT1000S). Samples were embedded in 5% low melting point agarose (TopVision, Thermo Scientific) on a 4% agar platform (NuSieve 3:1 Agarose, Lonza) and secured to the specimen disk using cyanoacrylate adhesive. Sectioning was performed in cutting solution (H+GPSA medium) using a sapphire blade (1.5-inch, Ted Pella, Inc.) advanced at 0.75 mm/s and oscillating at 60 Hz, with the chamber cooled to nominally –3 °C. Slices were 300 µm thick. However, rare loosely textured tumors were difficult to section and could be up to 2 mm thick.

Slices were placed on 30 mm-diameter cell culture inserts (0.4 µm pores, Millicell PICM0RG50) pre-coated to facilitate adhesion with a thin layer of growth factor-reduced, phenol red-free, Matrigel (Corning). Inserts were placed in 6-well plates containing 1.5 mL of GBS (Glioma Brain Slice) medium (modified from the medium described in Jacob et al., 2020 [42]): BrainPhys Neuronal Medium (StemCell Technologies) supplemented with: 1x GlutaMax (Thermo Fisher), 1x N2 supplement (Thermo Fisher), 1x B27 without vitamin A supplement (Thermo Fisher), 1x non-essential amino acids (NEAAs) supplement (Thermo Fisher), 1x Penicillin-Streptomycin (Thermo Fisher), 1x 2-mercaptoethanol (Thermo Fisher), and 2.5 µg/mL human insulin (Sigma). Cultures were maintained at 37 °C in a 5% CO_2_, 4% O_2_ environment for over 28 days. Media was fully replaced every 48-72 hours.

### Hematoxylin and Eosin (H&E) staining

Samples were fixed in 4% paraformaldehyde (PFA) in phosphate-buffered saline (PBS) for 30 min at RT, washed 3x in tris-buffered saline (TBS), then embedded in paraffin using a Tissue-*Tek* TEC Tissue Embedding Station (*SAKURA*), sectioned at 5 μm thickness, and positioned on positively-charged glass slides (Thermo Fisher). The slides were subsequently deparaffinized, rehydrated and stained on a Tissue-*Tek* VIP *Vacuum Infiltration Processor (SAKURA)* with Modified Mayer’s Hematoxylin and Eosin Y Stain (America MasterTech Scientific) on a H&E Auto Stainer *(Prisma Plus Auto Stainer, SAKURA)* according to standard laboratory procedures. Tissue sections were evaluated by a neuropathologist and imaged using a Hamamatsu NanoZoomer 2.0 HT slide scanner.

### Immunofluorescence staining

Intact tumor samples were carefully removed from the Millicell culture insert, fixed in 4% PFA for 30 min at RT and washed 3x in TBS. For frozen sectioning, samples were equilibrated in 30% sucrose overnight at 4 °C, embedded in Optimal Cutting Temperature (OCT) compound, and frozen on dry ice. Blocks were sectioned at 10-20 µm using a Tissue-*Tek* Cryo3 flex cryostat (*SAKURA*) and mounted on charged glass slides (Thermo Fisher). For immunostaining, frozen sections were thawed at RT for at least 10 min.

Fixed whole mount tumor samples (nominally 300 µm) or cryosections (10-20 µm) were blocked with 1x CAS-Block (Invitrogen, Thermo Fisher) for 1 hour at RT, then incubated with primary antibodies in blocking solution overnight at 4 °C, followed by secondary antibodies in blocking solution overnight at 4 °C. After washing, samples were mounted in Prolong Glass (Invitrogen) and stored in the dark at RT for at least 24 hrs before imaging.

### Confocal microscopy and image processing

Confocal imaging was performed at the City of Hope Light Microscopy and Digital Imaging core using Zeiss LSM880 or LSM900 microscopes. Optical slices (z-stacks) and tiling were configured based on the sample size. Images were processed using Zeiss ZEN Black and ZEN Blue software.

### PhenoCycler (CODEX) multiplexed imaging (Akoya Biosciences)

A panel of 21 CODEX antibodies was employed. Barcoded antibodies were purchased from Akoya, or carrier-free antibodies were purchased and barcode conjugated by Akoya. Details on these and other antibodies are provided in **Supplemental Table S3.**

PhenoCycler (CODEX) immunostaining and imaging were conducted according to the manufacturer’s instruction (CODEX User Manual – Rev C). Formalin-fixed, paraffin-embedded (FFPE) blocks were sectioned (10 µm) and placed onto glass coverslips (#1½; Electron Microscopy Sciences) pre-coated with 0.1% (w/v) poly-L-lysine (Sigma-Aldrich). Sections were baked at 60 °C for 25 min, and then deparaffinized in xylene, rehydrated through an ethanol series, and rinsed in ddH_2_O. Antigen retrieval was performed by incubating sections in 1x citrate buffer (Sigma-Aldrich, pH 6.0) in water for 20 min at 100 °C under pressure using a TintoRetriever pressure cooker (1709641; BIO SB Inc., NC). Sections were washed in hydration buffer (Akoya), then staining buffer (Akoya), and then stained with 190 µL of the 21-antibody cocktail in CODEX blocking buffer (Akoya) for 3 hours at RT in a humidified chamber. Imaging was conducted using a Zeiss Observer 7/PhenoCycler (CODEX) instrument with CODEX Instrument Manager (CIM) software. Raw data were analyzed with CODEX Multiplex Analysis Viewer software.

### RNA extraction and library preparation for bulk RNA-sequencing

For sample PBT293, RNA extraction was performed using the Qiagen RNeasy Mini Kit (Qiagen, Cat No. 74104) according to manufacturer’s instructions. RNA Quantification was conducted using the Qubit RNA HS Assay Kit (Fisher Scientific, Cat No. Q32852) and the Nanodrop One spectrophotometer (Fisher Scientific). Samples with adequate yield were submitted to the TGen Collaborative Sequencing Center and RNA-seq libraries were prepared using the Kapa RNA HyperPrep Kit with RiboErase (Roche, Cat No. KK8560). Libraries were sequenced on the Illumina NovaSeq 6000 platform with 2×100bp paired-end reads. For samples PBT308, PBT309 and PBT313, RNA was extracted using Qiagen RNeasy Mini kit (Qiagen), following manufacturer’s instructions. RNA quality and quantity were assessed using BioAnalyser Systems Tapestation 4200 (Agilent Technologies). Sample pairs that passed the RNA Integrity Number (RIN) evaluation were selected for sequencing library preparation using the KAPA RNA HyperPrep Kit with RiboErase (Roche). Sequencing of prepared RNA from parental GBM tumor and day 28 GOSC samples was performed on the Illumina NovaSeq 6000 using the S4 Reagent Kit v1.5 (Illumina) in paired-end mode of 2×151 cycles. The base calling was conducted using the Real-time Analysis (RTA) software v.3.4.4.

### Transcriptome analysis using bulk RNA-sequencing

Sequencing was performed at the COH Integrated Genomics Core or at the TGEN Clinical Genomics and Therapeutics Division. Sequence reads passing quality control were preprocessed for adapter and polyA removal using Trimmomatic v0.39 [43] and fastp v.0.23.2 [44]. Reads were aligned to human reference genome hg38, and indexed with gene and transcript annotations using STAR v.2.7.9a [45]. Strand-specific paired-end fragment counts for RefGene-annotated genes were obtained via featureCounts in the Subread package v.2.0.3 [46].

To enhance the accuracy of the gene expression estimates, 15,235 genes with counts per million (CPM) > 1 in at least four samples were selected for downstream analysis.

The Bioconductor package edgeR v.3.32.1 [47] with a design matrix based on the general linear model quasi-likelihood F-tests was used to evaluate differential gene expression between day 28 GOSCs and day 0 parental tumor pairs. For the volcano plot shown in Figure 2E, raw p-values were adjusted using the method of Benjamini and Hochberg for multiple-testing correction [48] to derive the FDRs (adjusted p-values) using 0.05 as the significance threshold. Based on the edgeR output, a FDR of 0.05 corresponds to a raw p-value of approximately 0.00178 in our case. This is the y-axis cutoff between colored and grey dots shown in the volcano plot: –log_10_(0.00178) = ∼2.75).

Pre-ranked Gene Set Enrichment Analysis (GSEA) [49] was conducted for functional enrichment analysis using gene sets from the human Molecular Signatures Database (MSigDB) v2024.1 [50]. Genes were ranked by logFC combined with –log_10_ of the p-value.

Additionally, Gene Set Variation Analysis was performed on selected core hallmark signatures using the Bioconductor package GSVA v.1.48.3 [51].

Principal Component Analysis was conducted using the R Stats package v.4.3.0 (R core team, 2025).

Cell type proportions were estimated with GBMDeconvoluteR [10] using the relative abundance of transcripts in units of counts per million (cpm) [52]. Marker genes for different GBM cell subtypes and for immune cells were sourced from two GBM single-cell reference datasets [10, 53].

### Analysis of public gene expression data

We used data from the Cancer Genome Atlas (TCGA) for glioblastoma (GBM), breast invasive carcinoma (BRCA), and pancreatic adenocarcinoma (PAAD), and the NCI Genomic Data Commons (GDC) for pancreas cancer organoid profiling (ORGANOID-PANCREATIC). RNAseq-STAR-Counts and phenotype files were obtained via UCSC Xena Functional Genomics Explorer [54].

The dataset included 157 primary GBM tumors (including 7 carrying IDH1-R132 mutations), 150 BRCA samples, 143 PAAD samples classified as ductal and lobular neoplasms, and 44 organoid samples from confirmed pancreatic ductal adenocarcinoma (PDAC) [55].

Gene identifiers (Ensembl IDs) were converted to gene symbols using the Bioconductor annotation package org.Hs.eg.db (DOI: 10.18129/B9.bioc.org.Hs.eg.db). A total of 22,426 annotated genes were analyzed. Batch effects across TCGA, GDC, and City of Hope (COH) samples were corrected using ComBat-seq in the Bioconductor package sva [56, 57]. Adjusted counts were normalized for library sizes. Multi-dimensional scaling (MDS) plots were generated using the Bioconductor package edgeR v.3.32.1 [47].

### CAR T cell generation and CAR T cell co-culture killing assay

Blood products were obtained under protocols approved by the City of Hope Internal Review Board from healthy donors. Peripheral blood mononuclear cells (PBMCs) were isolated by density gradient centrifugation over Ficoll-Paque (GE Healthcare, Little Chalfont, UK), followed by sequential rounds of CliniMACS/AutoMACS depletion to enrich and isolate the naïve/memory T cell (Tn/mem) population, as described previously [38].

Tn/mem cells were stimulated with Dynabeads Human T-Expander CD3/CD28 (Invitrogen) based on 2:1 bead/T cell ratio and transduced with lentiviral vectors encoding CAR constructs. T cells were cultured in X-Vivo-15/10% FBS media with 50 U/mL IL-2 and 0.5 ng/mL IL-15. One half the medium was changed every other day. After 7 days, the beads were removed, and T cells were counted using a Cellometer K2 cell counter before expansion. The percentage of CAR-expressing T cells was determined by flow cytometry (Attune NxT Flow Cytometer, Thermo Fisher) using a truncated CD19 construct co-expressed with the CAR. Data were analyzed by FCSexpress (De Novo Software).

E:T (Effector:Target) ratios were fixed at 1:1 for all CAR T cell killing experiments, with the appropriate numbers of effector cells required determined from the total number of DAPI+ cells in each GOSC. To calculate this, we first estimated the total number of cells in each GOSC. As no significant change of cell density was observed in GOSCs compared to the parental tumor samples (**Supplemental Figure S3B**), we determined densities of DAPI+ cells in representative 300 µm-thick slices of parental tumors by confocal imaging. We then used Imaris software (v9; Oxford Instruments) to count the total number of DAPI+ cells (*X*) and measure the cross-sectional area (*S*_0_) of representative parental tumor sections. Then, with *H* being the total height of the parental tumor section, the cell density (*D*) per µm^3^ in the parental tumor sample section was:

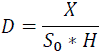

Next, to apply this number to each tumor’s GOSCs, we brightfield imaged each living GOSC on its culture insert (before applying CAR T cells) and measured its cross-sectional area (*S*_n_) with Fuji/ImageJ. Since each GOSC was initially 300 µm thick, the total number of cells in each GOSC (*N*_GOSC_) was used to calculate the number of CAR T cells to apply for a given E:T ratio:

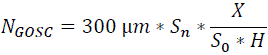

The computed numbers of CAR T cells or Mock T cells to be used were centrifuged, resuspended in 5 µL of X-Vivo medium, and gently added to the upper surface of a GOSC on its culture insert. Co-culture assays were conducted at 37 °C in the 5% CO_2_, 4% O_2_ environment. Seven days after T cells were added to the GOSCs, cultures were fixed in 4% PFA for 30 min at RT. Immunofluorescence staining for CD3 (T cells), cleaved caspase 3 (CC3, cell apoptosis), and IL13Rα2, and streptavidin:fluorophore staining for CLTX:biotin, were conducted as described above.

### Image quantification and statistical analysis

All CODEX and immunofluorescence images were quantified using QuPath v0.7.0 [58]. Three representative sample regions were selected for annotation, and cell numbers were determined from DAPI staining. To quantify markers expressed in cytoplasm, we used the pixel classification function to train QuPath and then measured the percentage of positive area (area of positive expression/area of total slice) automatically. To quantify markers expressed for characterization of tumors, including SOX2 and OLIG2, immune cell markers CD3, CD68 and CD163, and cell apoptosis marker cleaved caspase 3 (CC3), we used the object classification function to train QuPath and measured the percentage of positive cells (number of immuno-positive cells/number of DAPI+ nuclei). In the CAR T cell killing assays, it was more accurate to count the number of CD3+ cells manually. All quantifications were examined for statistical significance by two-way ANOVA with multiple comparison corrections (Prism v10; GraphPad). All data are presented as mean ± SD, with statistical significance noted in each figure as: ** (p<0.01), * (p<0.05) or ns (p>0.05, not statistically significant).

## RESULTS

### Histopathological features of human high-grade glioma (HGG) are maintained in our long-term organotypic slice culture system

Freshly resected patient tumor samples (patient information is summarized in **Supplemental Table S1**) were sliced at 300 µm using a vibratome to generate ex vivo organotypic HGG cultures with the smallest possible disturbance to tissue architecture. Tumor slices were placed on tissue culture inserts and grown at the air-liquid interface in a defined serum– and growth factor-free BrainPhys-based medium under conditions approximating those found in vivo (2.5 mM glucose, 4% O_2_) [35, 59] for at least 28 days **(Figure 1A)**. Eleven patient samples were of sufficient quality when received from surgery to be cultured.

**Figure 1.**
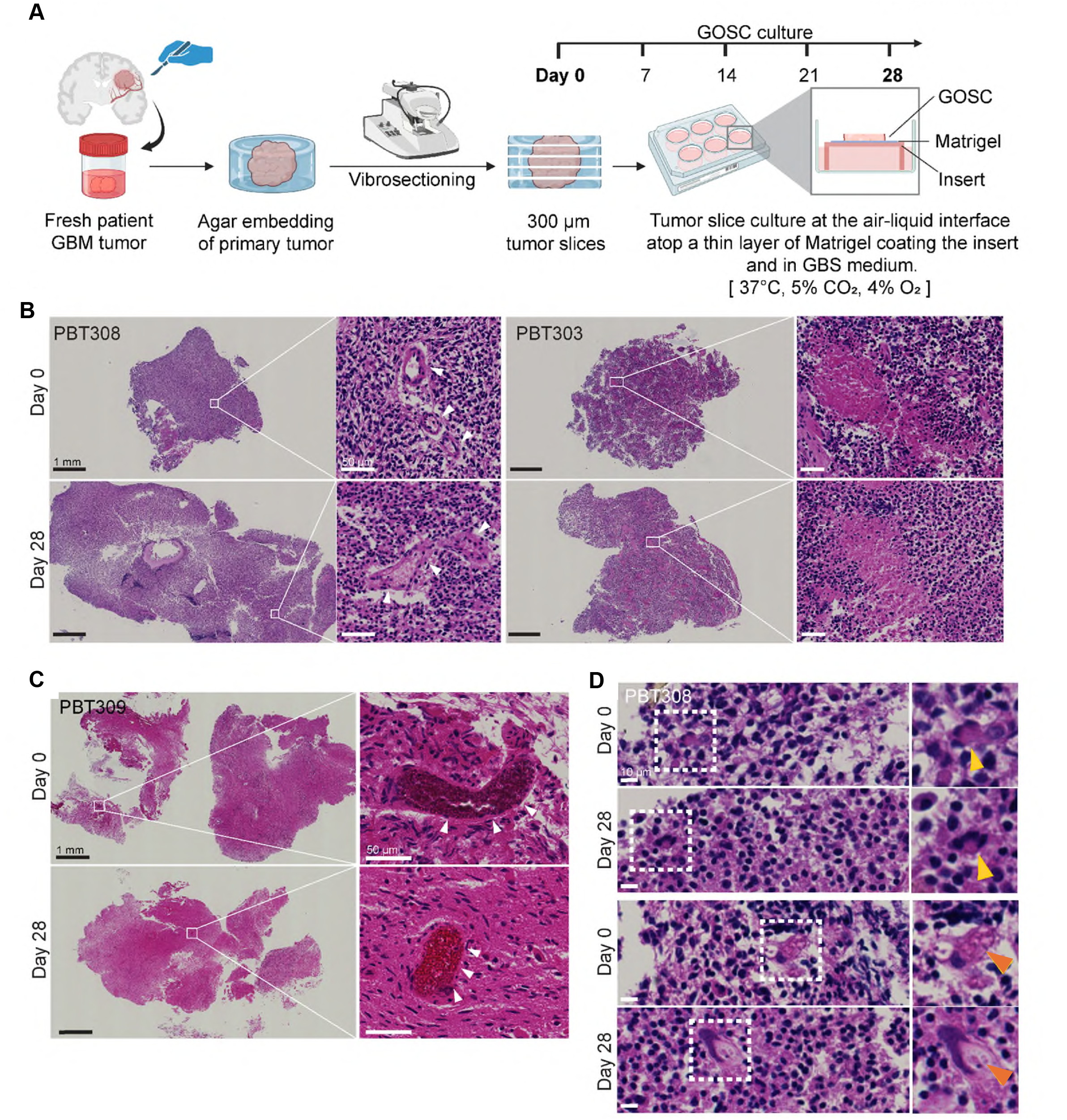
Organotypic GOSCs preserve histopathological features of parental patient HGG tumors for 28 days. **(A)** Schematic of the procedure to generate 300 µm-thick GOSCs from patient tumor samples and their organotypic culture in GBS medium (see Methods). Created in BioRender. **(B)** GOSCs retained tumor vasculature (PBT308) and areas of pseudopalisading necrosis (PBT303) characteristic of CNS WHO grade 4 gliomas (GBM). Shown are H&E-stained images of PBT308 day 0 parental tumors (top left) and PBT303 (top right), paired with and their corresponding day 28 GOSCs (bottom left and bottom right). Higher magnification details in PBT308 illustrate tumor vasculature, and in PBT303 illustrate necrotic regions with pseudopalisading necrosis. Black scale bars in full images = 1 mm; white scale bars in insets = 50 µm. See **Supplemental Figure S1A** for additional examples of retained tumor structures. **(C)** Erythrocytes were retained in tumor vasculature. H&E staining of tumor PBT309 parental day 0 (top) and corresponding day 28 GOSC (bottom); white arrowheads mark vasculature walls enclosing erythrocytes. Scale bars as in (B). **(D)** GOSCs retain cells with preserved cytomorphological features observed in the original WHO grade 4 tumor. Shown are H&E staining of two regions of PBT308 day 0 parental tumors and corresponding day 28 GOSCs; insets show the areas indicated. There is diffuse nuclear hyperchromatic staining associated with marked nuclear pleomorphism, multinucleation and focally abundant gemistocytic cytoplasm with focal vacuolization. Arrow heads mark multinucleation (yellow) and vacuolated gemistocytic cytoplasm (orange). Scale bar on left = 10 µm.

**Figure 2.**
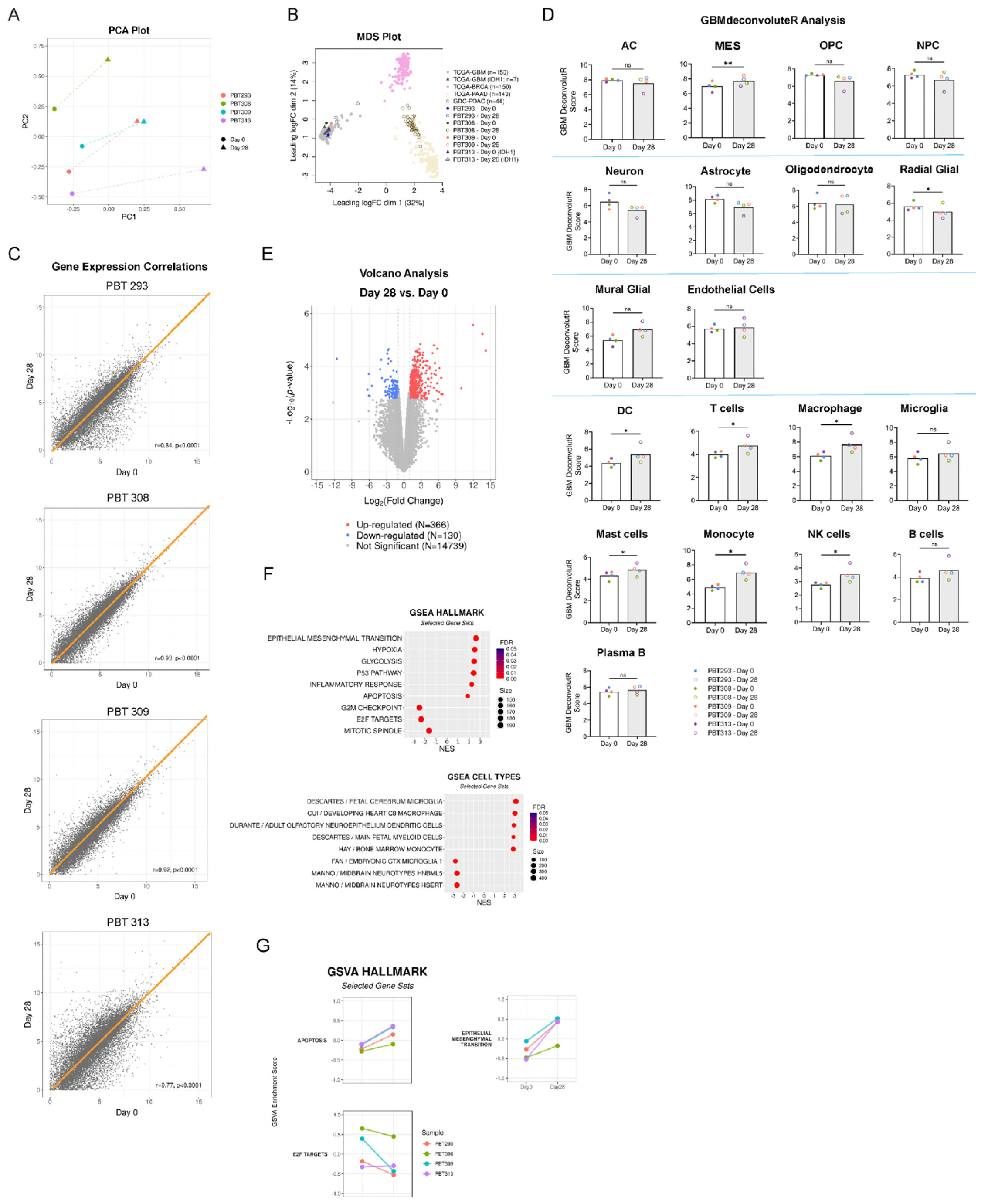
Bulk RNA-seq expression profiling illustrating clustering and correlations of day 0 parental HGG tumors and their corresponding day 28 GOSCs. **(A)** Principal Component Analysis (PCA) of transcriptomic profiles from paired patient-derived samples. PCA was performed on log2-transformed counts per million (CPM) data to assess the global variance between parental tumor samples and GOSC samples over time. Each color represents a unique patient-derived sample, while shapes indicate the time point in culture: solid circles represent day 0 and triangles represent day 28. Dashed lines connect paired samples from the same subject. PC1 and PC2 represent the first and second principal components, respectively. **(B)** Gene expression multi-dimensional scaling (MDS) plots. Shown are clustered bulk RNA-seq data from parental and corresponding day 20 GOSCs for three IDH1-wildtype tumors (PBT293, PBT308, PBT309) and one IDH1-mutated tumor (PBT313) (comparable data from IDH-mutated PBT261 and PBT289 were not available). These are superimposed on clustered data for primary tumors drawn from public data sets: glioblastoma, breast cancer and pancreatic cancer. Sources: GBM, TCGA (n = 157) (157 primary tumors, including 7 carrying IDH1 mutations); BRCA (breast cancer), TCGA (n = 150); PAAD (pancreatic cancer) TCGA (n = 143) (primary tumors “ductal and lobular neoplasms” to match the GDC organoid dataset); GDC organoid-pancreatic (n = 44) (derived from PDAC; with “disease type unknown” removed). For project information, go to https://portal.gdc.cancer.gov/ and input above highlighted project names in the search bar. **(C)** Pearson correlations of gene expression determined from bulk RNA-seq illustrating correlations between day 0 parental tumors and corresponding day 28 GOSCs for three IDH1 wildtype patient tumors (PBT293, PBT308, PBT309) all with r > 0.9 and one IDH1-mutant tumor (PBT313) with r = 0.77. **(D)** GBMdeconvoluteR analysis of HGG cell type marker gene expression (as indicated) of day 0 parental tumors and corresponding day 28 GOSCs. The presence of most cells determining tumor structure, including neural stem cell-derived lineages, were stable: astrocytic cells (AC), oligodendrocyte precursor cells (OPC), neural progenitor cells (NPC), and mature neurons, astrocytes and oligodendrocytes. Vascular cells shaping tumor architecture (mural glial, endothelial) were stable, as well as microglia (resident macrophages). Radial glial cells decreased, perhaps reflecting a transition towards the MES tumor cell subtype. The presence of cells comprising the tumor microenvironment, often innate and adaptive immune cells invading active tumors, increased somewhat over time: dendritic cells (DC), T cells, macrophages, mast cells, monocytes, NK cells. A component of adaptive immunity, B-lineage cells (plasma B cells), remained stable. *Note that for GBMdeconvoluteR analyses, valid quantitative comparisons can only be made within a cell type (day 0 vs. day 28 in this case) but not between data for different cell types*. **(E)** Differential gene expression analyses between day 0 parental tumors and corresponding day 28 GOSCs. Shown is a volcano plot of differentially expressed genes: 496 of 15235 total genes (red, significantly up-regulated; blue, significantly down-regulated, FDR < 0.05). Expression of 3.3% of total genes significantly changed over time. **(F)** Bubble plots from gene set enrichment analysis (GSEA) using the Hallmark and Cell Type molecular signature databases, with significantly down– and up-regulated pathways shown. Within the GSEA Hallmark series, five of six up-regulated components were related to overall tumor status: epithelial-mesenchymal transition, hypoxia, glycolysis, inflammatory response, apoptosis. E2F targets (related to the cell cycle, DNA replication, and mitosis) were downregulated. **(G)** Selected hallmark gene set enrichment signatures over 28 days of culture. Enrichment scores were calculated using Gene Set Variation Analysis (GSVA) for selected Molecular Signatures Database (MSigDB) Hallmark pathways. Each row represents a distinct biological pathway, with y-axes manually fixed from –1 to 1 to facilitate direct magnitude comparison across gene sets. Data points represent individual sample pairs at day 0 and day 28, with colors corresponding to specific samples. Lines connect paired longitudinal samples.

We first compared the histopathological features of uncultured parental HGG tumors (day 0) to their matched cultured day 28 GOSCs using hematoxylin and eosin (H&E) staining on formalin-fixed, paraffin-embedded (FFPE) sections. On day 28, viable cells were present in all GOSC derived from seven patients examined with H&E staining (**Figure 1B-1C, Supplemental Figure S1A)**. Importantly, tumor vasculature, including venules and capillaries, was maintained in the GOSCs **(Figure 1B-1C, Supplemental Figure S1A)**. Interestingly, erythrocytes remained visible in the GOSC tumor vasculature even after 28 days **(Figure 1C)**.

Pseudopalisading necrosis, a hallmark of WHO grade 4 glioma [6], was observed in some pairs of parental tumors and their corresponding GOSCs **(Figure 1B: PBT303)**. GOSCs exhibited other diagnostic features of HGGs, including mitotically active, enlarged, irregular neoplastic nuclei on a fibrillary background (**Figure 1B, Supplemental Figure S1A**).

GOSCs retain cells with preserved cytomorphological features observed in the original WHO grade 4 tumor **(Figure 1D)**. Shown are H&E staining of three regions of PBT308 day 0 parental tumors and corresponding day 28 GOSCs, with insets show the areas indicated. There is diffuse nuclear hyperchromatic associated with marked nuclear pleomorphism, multinucleation and focally abundant gemistocytic cytoplasm with focal vacuolization. Arrow heads mark multinucleation (top image pair) and vacuolated gemistocytic cytoplasm (bottom image pair).

Non-tumor regions, including white and grey matter at the tumor edges and infiltrating areas, were also preserved **(Supplemental Figure S1B-S1C).** Together, these results demonstrate that our GOSC platform maintains key histopathological features, macrostructures, and peritumoral regions of patients’ HGG tumors for at least 28 days.

### Gene expression signatures of parental tumors were retained in day 28 GOSCs

To evaluate preservation of HGG gene signatures, we performed bulk RNA expression profiling on day 28 GOSCs for four patients (PBT293, PBT308, PBT309, and PBT313) and compared these results to those of their cognate parental tumors.

We used principal component analysis (PCA) analysis to test for potential non-biological contributions to differences between the cultured tumor samples (**Figure 2A**). The maintained positions of each day 0-day 28 tumor pair relative to the others indicate that biological identity is the primary driver of transcriptomic variance, confirming the absence of significant batch effects or substantial culture-induced drift over the 28-day period.

To visualize the similarity or dissimilarity between samples, we performed a multidimensional scaling (MDS) plot analysis comparing our eight samples with other bulk RNAseq samples from the Cancer Genome Atlas (TCGA) (**Figure 2B**): glioblastoma (n = 150; grey), GBM with IDH1 mutation (n = 7; dark grey), breast cancer (n = 150; lavender pink), pancreatic cancer (n = 143; pale green), and pancreatic cancer organoids (n = 44; open olive-green).

As expected, cancers from different organs (brain, breast or pancreas) were clustered separately, with our day 0 parental GBM samples positioned within the TCGA glioblastoma cluster. Our day 28 GOSCs samples were also positioned within the main GBM cluster, indicating stability of gene expression patterns. However, notably, the day 28 GOSC of PBT313 bearing an IDH1 mutation (open triangle), fell to the margin while still within the overall GBM cluster, suggesting possible influence of the IDH1 mutation on progression of the GOSC. This will be explored in the future. The cultured pancreatic cancer organoids (open olive-green) were also slightly shifted from the primary PDAC tumor cluster (pale green), suggesting that MDS shifts within cluster boundaries may reflect modest transcriptional changes during the culture process.

Next, Pearson correlation analyses, used to examine stability of gene expression between parental tumors and day 28 GOSCs (**Figure 2C**), revealed high correlations for patients PBT293, PBT308, and PBT309, with R values of 0.84, 0.93, and 0.92, respectively. Interestingly, for PBT313 carrying the IDH mutation, and consistent with the result in the MDS plot, the R value was the lowest at 0.77. This may reflect the reduced survival of IDH-mutant glioma cells in culture [60] or other factors. Nevertheless, as R values between 0.7 to 1.0 are indicative of strong positive correlation, these results indicate that day 28 GOSCs largely preserved the gene expression profiles of their parental tumors.

### GOSCs retain the full complement of cell types in parental HGG tumors as indicated by GBMdeconvoluteR analysis

Within HGGs, non-tumor cellular components of the tumor microenvironment (TME) significantly impact tumor development and progression and are a driving force in therapy resistance [50]. We therefore examined whether GOSCs maintain the TME cells resident in parental HGG tumors. To determine the types of cells present in day 28 GOSCs, we employed the web application tool GBMdeconvoluteR to quantify the proportions of GBM cell types in parental and day 28 GOSCs based on bulk RNA-sequencing data [10, 61] (**Figure 2D**).

We first compared the relative abundance of four neoplastic cell types: neural-progenitor-like (NPC), oligodendrocyte-progenitor-like (OPC), astrocyte-like (AC), and mesenchymal-like (MES) cells, and observed a small increase in the abundance of MES-type cells but no changes in other cell types. These results suggest the preservation of NPC, OPC and AC cells, and raise the possibility of some epithelial-to-mesenchymal transition (EMT) in day 28 GOSCs. Next, we examined the abundance of brain neural-lineage cells (astrocytes, oligodendrocytes, neurons, and radial glial cells), vasculature cells (mural and endothelial cells), and immune cells (B cells, T cells, NK cells, microglia, tumor associated macrophages, monocytes, mast cells, and dendritic cells). Results showed the preservation of astrocytes, oligodendrocytes, neurons, mural, and endothelial cells, microglia, B cells, and plasma cells. However, the abundance of radial glial cells (presumptive neural stem cells) decreased, and the abundance of dendritic cells, macrophages and T cells increased. We concluded that based on GBMdeconvoluteR gene expression analysis, all cell types were retained by day 28 GOSCs, with small changes in the abundance of some immune cells relative to parental tumors.

### Gene pathway analysis of day 28 GOSCs indicated an increase in the relative presence of inflammatory cells as compared to day 0 GOSCs

A volcano plot analysis of a total of 15,235 genes in paired day 0 *vs*. day 28 GOSCs revealed that 96.7% (14,739 genes) were not significantly changed (FDR > 0.05) over this time **(Figure 2E)**. Among the 3.3% genes that were statistically different (FDR < 0.05), 130 genes were significantly down-regulated, and 366 genes were up-regulated.

To gain insight into the functional implications of these changes, we conducted gene set enrichment analysis (GSEA) [49, 62] using the Hallmark (**Figure 2F, top**) and Cell Type (**Figure 2F, bottom**) molecular signature databases to identify pathways and cell phenotypes impacted by GOSC culture. Most up-regulated pathways in day 28 GOSCs identified by the Hallmark database were consistent with an enhanced tumorigenic phenotype, including genes involved in EMT, responses to hypoxia, increases in glycolysis, the p53 pathway, inflammatory responses, as well as apoptosis (**Figure 2F, top**). The up-regulated EMT and inflammatory response genes are consistent with the conclusions of the GBMdeconvoluteR analysis (**Figure 2D**). Down-regulated gene pathways include those involved in G2M checkpoint, E2F targets, and mitotic spindles (**Figure 2F, top**).

Similarly, GSEA cell type analysis indicated that many of the up-regulated gene sets identified monocyte, macrophage, and microglia lineages (**Figure 2F, bottom**), again consistent with GBMdeconvoluteR analysis (**Figure 2D**). Conversely, down-regulated gene sets identified embryonic microglia and midbrain neurotypes (**Figure 2F, bottom**). Taken together, these results suggest that certain immune cell types, including microglia, are enriched in day 28 GOSCs as compared to parental tumors.

To characterize the transcriptomic evolution of the samples during culture, we performed Gene Set Variation Analysis (GSVA) (**Figure 2G**) on selected core hallmark signatures for proliferation (E2F TARGETS), apoptosis (APOPTOSIS), and migration (EPITHELIAL MESENCHYMAL TRANSITION). Quantitative comparison of enrichment scores confirmed significant longitudinal shifts in these core biological programs while preserving overall tumor cell types and architecture. As visualized in the paired dot plots, we observed highly consistent directions of change across patient-derived GOSCs, mirroring the significant up– and down-regulation trends (FDR < 0.05) identified in our GSEA analysis. This alignment between GSEA and sample-specific GSVA analyses underscores a reproducible evolution in immune cell states over the 28-day period.

### Immunofluorescence staining of cell and matrix markers confirmed preservation of all cell types in day 28 GOSCs

#### Immunotherapy target antigens

Expression of three well-known tumor antigen targets for CAR T cell immunotherapies, EGFR, IL13Rα2, and HER2, was evaluated in a group of four day 28 GOSCs and in their respective parental HGG tumors (**Figure 3A, Supplemental Figure S2A-S2C**), without statistically significant changes in overall cell density (**Supplemental Figure S2D**) or in numbers of antigen-expressing cells (**Figure 2F**). Stability of the presence or absence of parental tumor antigens in the GOSCs will be important for future evaluations of immunotherapies using this GOSC platform, as described below.

**Figure 3.**
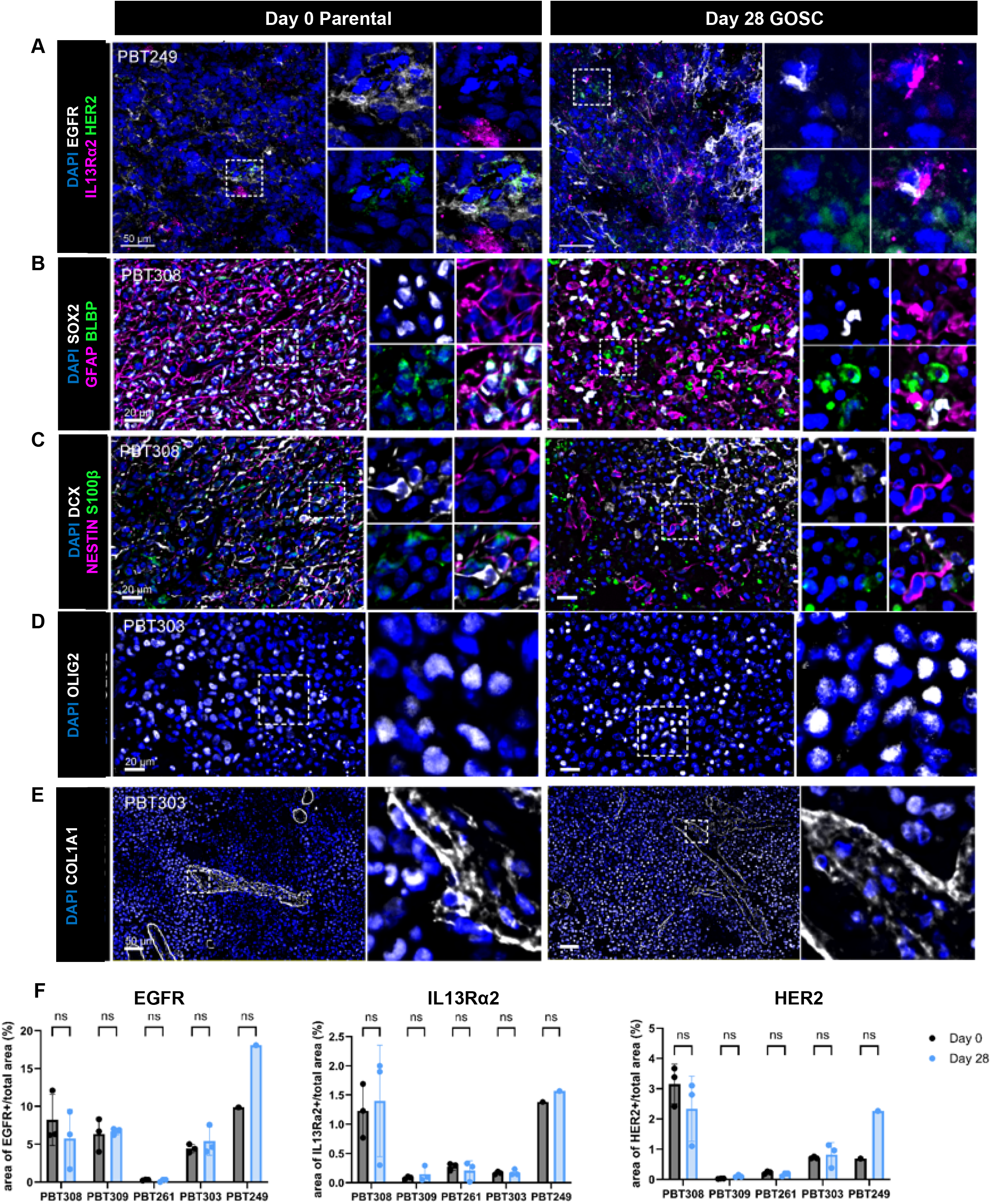
Day 28 GOSCs preserve cell populations of day 0 parental tumors as determined from immunostaining: CAR T cell tumor antigen targets, glioma stem/progenitor cells, neuronal lineage cells, and extracellular matrix components. **(A)** Immunostaining of day 0 and day 28 PBT249 tumor indicates maintenance of three tumor antigen immunotherapy targets: HER2 (green), IL13Rα2 (magenta), EGFR (white). Blue: DAPI stained nuclei. Additional examples are shown in **Supplemental Figure S2A**. Scale bar = 50 µm. **(B, C)** PhenoCycler (CODEX) immunostaining for markers of glioma stem/progenitor cells and of neuronal lineage cells in day 0 and day 28 PBT308 tumor. (B) Representative images of: BLBP (green), GFAP (magenta), SOX2 (white), and DAPI (blue). (C) Representative images of: S100β (green), NESTIN (magenta), DCX (white), and DAPI (blue). Quantified data are shown in **Supplemental Figure S4A, S4B**. Scale bar = 20 µm. **(D)** PhenoCycler (CODEX) immunostaining for oligodendrocyte marker OLIG2 (white) and nuclei (DAPI, blue) of day 0 and day 28 PBT303 tumor. Quantified data are shown in **Supplemental Figure S4C**. Scale bar = 20 µm. **(E)** PhenoCycler (CODEX) immunostaining for collagen marker COL1A1 (white) and nuclei (DAPI, blue) of day 0 and day 28 PBT303 tumor. Quantified data for COL1A1 expression are shown in **Supplemental Figure S4D.** Scale bar = 20 µm. **(F)** Quantification of immunostaining for the immunotherapy target antigens illustrated in **A** and **Supplemental Figure S2A** as percent of total area. Data are mean ± SD for three representative regions.

#### Glioma stem/progenitor cells

In human HGG, these cells contribute to tumor angiogenesis and therapy resistance [13, 22]. Thus, we stained for neural stem/progenitor and glioma stem cell markers NESTIN [63], BRAIN LIPID BINDING PROTEIN (BLBP) [63], and SOX2 [63]. All three markers were observed in GOSCs after 28 days in culture (**Figure 3B-3D, Supplemental Figure S3**). The immature neuron marker DOUBLECORTIN (DCX) [64], glia markers GFAP [65] and S100β [65], and oligodendrocytes marker OLIG2 [66] were also detected in GOSCs **(Figure 3B-3D, Supplemental Figure S3)**. No statistically significant changes of the expression of these markers between parental tumor samples and day 28 GOSCs were observed (**Supplemental Figure S4A-S4C**), indications of overall GOSC stability.

#### Extracellular matrix

In human HGG, the extracellular matrix protein COLLAGEN1A1 (COL1A1) plays a key role in tumor malignancy [67] by remodeling the tumor microenvironment, and supporting tumor proliferation, invasion and progression [67, 68]. We found that COL1A1 protein in day 28 GOSCs was maintained from the corresponding parental tumors (**Figure 3E**), without statistically significant changes (**Supplemental Figure S4D**).

#### Vascular cells

Hypervascularization is a hallmark feature of HGG [69], as abnormal and dysfunctional vasculature within HGGs contributing to the immunosuppressive microenvironment, tumor progression, and tumor angiogenesis [7, 13, 70]. We therefore examined four vasculature proteins (**Figure 4, Supplemental Figure S5**): endothelial cell markers von-Willebrand factor (vWF) and CD31 [71], endothelial cell tight junction marker ZO1 [72], and ensheathing smooth muscle actin (SMA) [73]. Expression patterns of all four markers in day 28 GOSCs indicated preserved tumor vasculature, as evidenced by similar levels of CD31, vWF and SMA expression in GOSCs and their corresponding parental tumors. Taken together, these results are consistent with the results of H&E staining (**Figure 1B-1C, Supplemental Figure S1**) and demonstrate that tumor vasculature is maintained in GOSCs at day 28.

**Figure 4.**
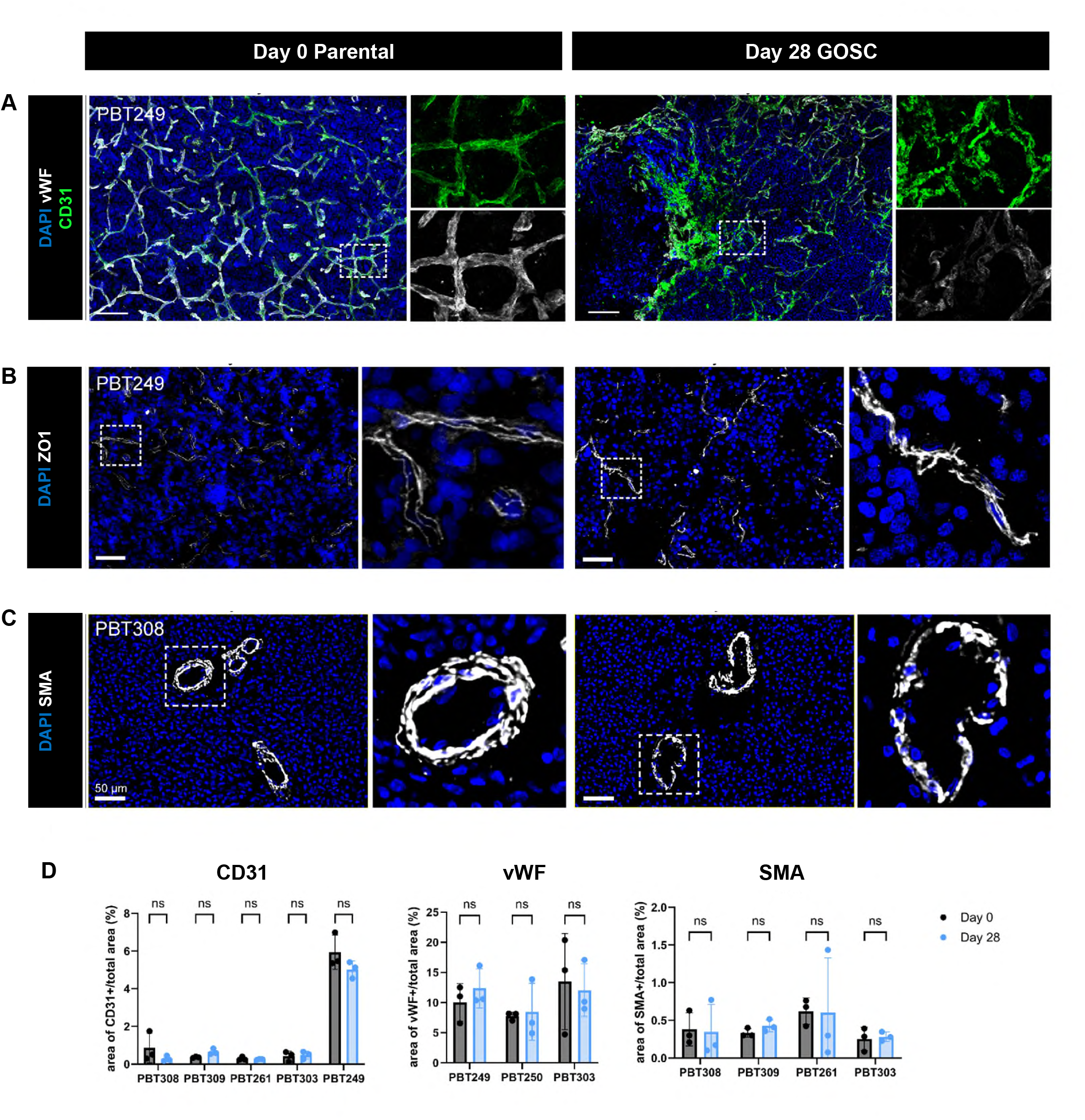
Day 28 GOSCs preserve vasculature of day 0 parental tumors. **(A, B)** Representative confocal images (z-axis projection) of nuclei (DAPI, blue) of day 28 and parental day 0 tumors, together with immunostaining for **(A)** vasculature markers CD31 (green), von Willebrand factor (vWF, white) (PBT249), **(B)** tight junction marker ZO1 (white) (PBT249). Scale bars = 100 µm for A, 50 µm for B **(C)** PhenoCycler (CODEX) immunostaining for muscle cell marker SMA (white) (PBT308). Scale bar = 50 µm. **(D)** No statistically significant differences were seen in quantified measures of CD31, von Willebrand factor, and SMA expression between day 28 GOSCs and day 0 parental tumor samples for the PBT tumor pairs indicated. Data (mean ± SD) are presented as area of positive expression (number of positive pixels, in units of µm^2^) divided by the total area (µm^2^). See **Supplemental Figure S5** for additional examples.

#### Immune cells

Multiple types of immune cells are present in human HGGs and their crosstalk with other cells and extracellular matrix components in the microenvironment play crucial roles in tumor development, progression, invasion, and responses to treatment [74]. Therefore, we examined (**Figure 5A, 5B, Supplemental Figure S6**) T cell marker CD3 [75], macrophage/microglia/monocyte markers CD14 [75] and CD68 [76], and M2-phenotype macrophage marker CD163 [75]. Cells positive for all these markers were detected in day 28 GOSCs. Quantification of CD3+ T cells and CD68+, CD163+, and CD14+ macrophage/microglia cells in day 28 GOSCs indicated that they were maintained at levels similar to those of their parental tumors **(Figure 5C)**.

**Figure 5.**
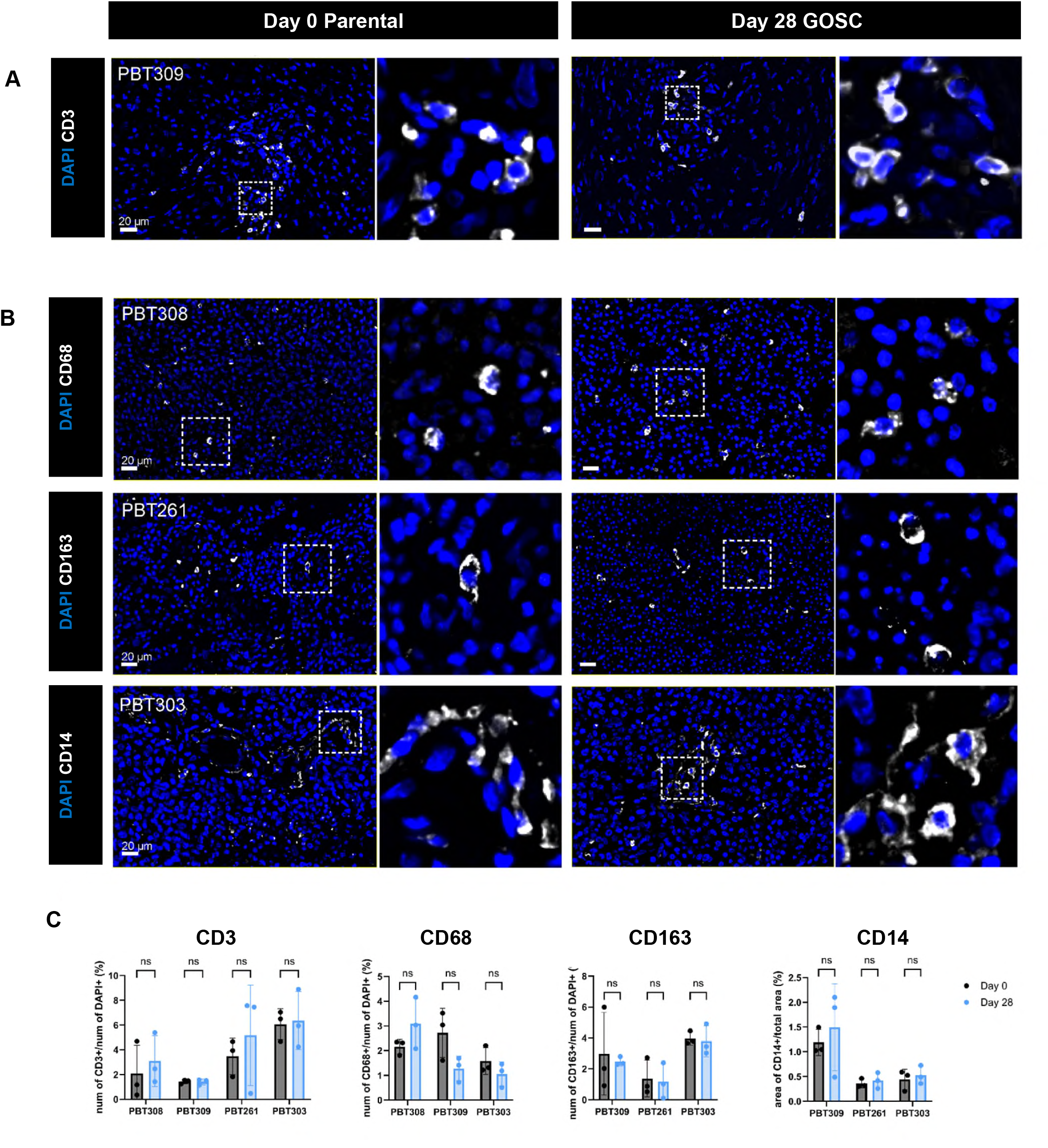
Day 28 GOSCs maintain immune cells of day 0 parental tumors. **(A, B)** Representative PhenoCycler (CODEX) images of nuclei (DAPI, blue) and **(A)** immunostaining for T cell marker CD3 (white) (PBT303), and **(B)** microglia/macrophage/monocytes markers (all white): CD68 (top), CD163 (middle), and CD14 (bottom) (PBT303). Scale bars = 20 µm. **(C)** Quantification of CD3, CD68, CD163 and CD14 on day 28 GOSCs and day 0 parental tumor samples for the PBT tumor pairs indicated. Data (mean ± SD) are presented as the number of positive cells divided by the total number of DAPI+ nuclei for CD3, CD68 and CD163, and as areas of positive expression (counted as numbers of pixels and presented as µm^2^) divided by the total area (µm^2^) for CD14. See **Supplemental Figure S6** for additional examples.

### Multiplex immunostaining indicates that the spatial arrangements of the multiple cell types comprising a tumor mass and shaping its architecture are preserved in day 28 GOSCs

In previous sections, we showed that examined individually, tumor antigen expressing cells (**Figure 3A, Supplemental Figure S2C**), glioma stem/progenitor cells (NESTIN+, SOX2+, BRAIN LIPID BINDING PROTEIN [BLBP]+) (**Figure 3B**), developing neuronal cells (DOUBLECORTIN [DCX]+) (**Figure 3C**), tumor cells or astrocytes (GFAP+, S100β+) (**Figure 3C, 3D**), oligodendrocyte lineage cells (OLIG2+) (**Figure 3D, Supplemental Figure S4**), endothelial (CD31+)/vasculature (von Willebrand factor)+/smooth-muscle-like (SMA+) cells (**Figure 4**) and immune cells in multiple lineages (CD14+, CD68+, CD163+, CD3+) (**Figure 5A, 5B**) were preset in day 28 GOSCs at levels similar to their parental tumors (**Figure 5C**). Overall, these observations suggest that proportions of the different cell types comprising a parental tumor mass are preserved in day 28 GOSCs.

We next used the multiplexing capability of PhenoCycler (CODEX) immunostaining to comprehensively visualize multiple cell lineages within a single tissue sections [77] and compare macro-organization of day 28 GOSCs to their parental tumors. An example is PBT309, for which **Supplemental Figures S7 and S8** show the GOSC and parental tumor, respectively (representative of comparisons additionally made for PBT261, PBT303 and PBT308). In these images, an area of vasculature and surrounding tumor in the multiplexed image (upper left) is highlighted and deconstructed into individual antigens using endothelial cells (CD31+) and smooth muscle actin cells (SMA+) as a vascular reference structure with its edges marked by the open arrows. The arrows are in registration around this vasculature and positioned based on positions of nuclei. We found that most strongly for PBT309 GOSC as well as its parental tumor, IBA1+, CD163+, CD68+, CD14+ cells of innate immunity, and CD3+ T cells of adaptive immunity, were clustered around vasculature, in particular the CD31+/SMA+ plexus just above and to the right of the main vessel. Extracellular matrix component collagen 1A1 (COL1A1) formed a tight band just outside the CD31+/SMA+ cells. GFAP+ astrocytic and tumor cells were specifically excluded from peri-vascular regions with a tightly defined edge and distributed throughout the tumor. Cells of NSC-derived lineage and their precursors, BLBP+ (radial glia), SOX2+ and OLIG2+ (stem cells), DCX+ (early migrating neurons), and S100β+ (astrocytes) were distributed through the tumor parenchyma. Curiously NESTIN+ cells (neuron progenitors) were present at higher density in the area around the main vessel as well as in the parenchyma. Taken together, these multiplex protein staining observations indicate that day 28 GOSCs preserve the complex architecture and spatial arrangements of cells within parental HGG tissues.

### GOSCs can be utilized as an ex vivo model to evaluate CAR T cell immunotherapy

To examine the potential utility of our GOSCs as a platform for modeling therapies for HGG, we assessed preservation of target specificity for CAR T cell treatments **(Figure 6A)**. Previously, we reported on chlorotoxin (CLTX)-directed CAR T cells, which exhibit anti-HGG activity and target tumors both in vitro and in vivo [37]. CLTX is a 36-amino acid peptide that binds to a broad spectrum of brain tumor cells but not adjacent or distant non-malignant cells [78].

**Figure 6.**
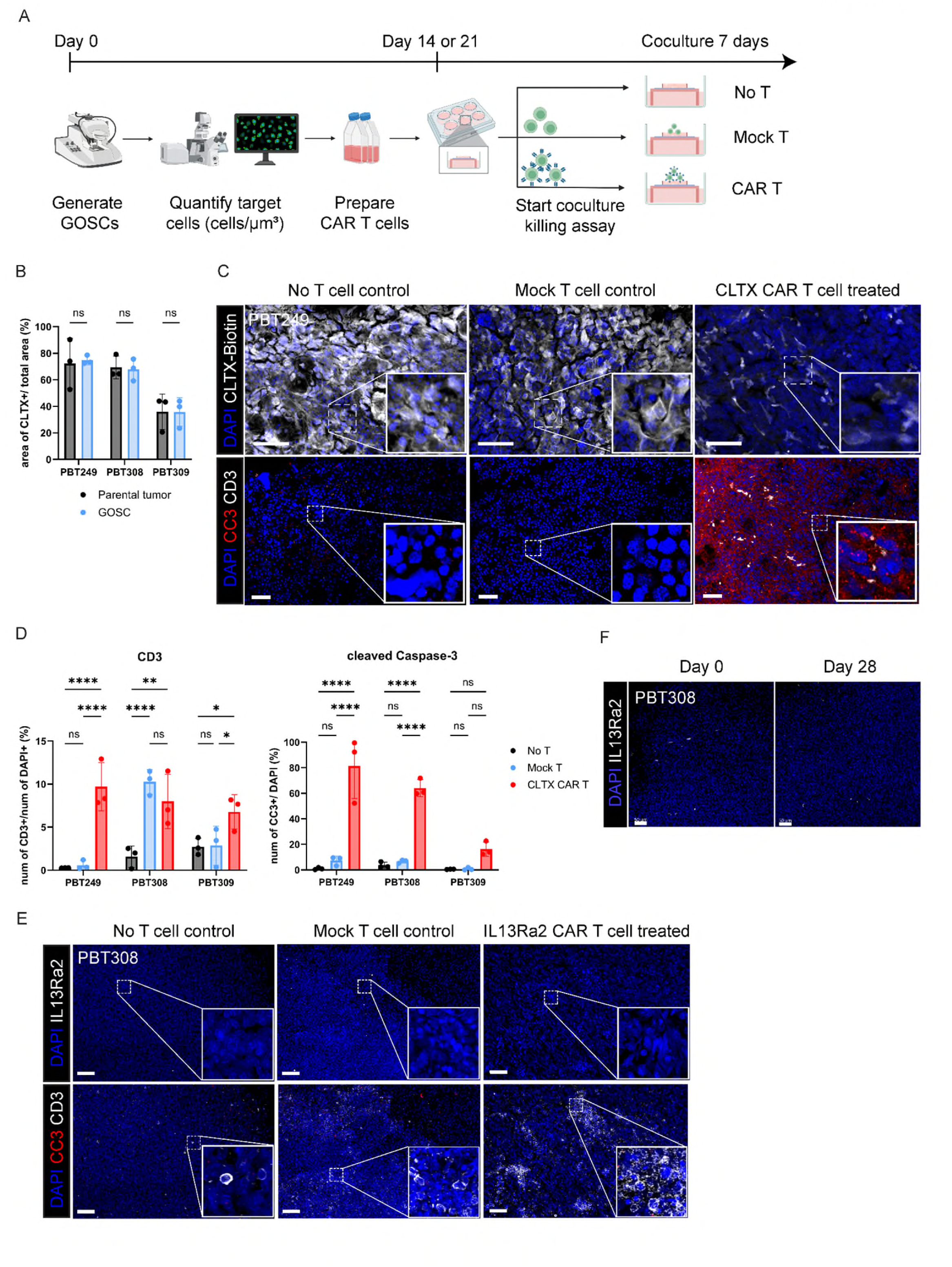
Evaluation of CAR T cell killing on GOSCs by co-culture killing assays. **(A)** Schematic depicting the CAR T cell co-culture killing assay on GOSCs. Conditions were: no T cells added control, mock T cell control, or CAR T cells (CLTX-CAR or IL13Rα2-CAR T cells). Created in BioRender. **(B)** Preservation of CLTX:biotin binding (visualized with streptavidin:AlexaFluor647) determined as area of streptavidin:AF647 fluorescence (number of positive pixels, in µm^2^) divided by the total area (µm^2^) for three parental and day 28 GOSC pairs (PBT249, PBT308, PBT309). **(C)** Specificity of CLTX-CAR T cell killing is maintained in day 28 GOSCs. Representative confocal images (z-axis projections) of streptavidin:AlexaFluor647 staining for CLTX:biotin (top, white), and immunostaining for CD3 (bottom, white) and CC3 (bottom, red) for PBT249 GOSCs after 7 days co-culture with either no T cells added, mock T cells, or CLTX-CAR T cells. E:T ratio = 1:1. DAPI (blue) shows nuclei. Scale bar = 50 µm. **(D)** Quantification (mean ± SD) of CLTX-CAR T cell infiltration (CD3+, left) and killing (CC3+, right) (red bars) for three GOSCs (PBT249 [panel C], PBT308, PBT309) in 7-day CLTX-CAR T cell co-culture killing assays. Data are presented as percent of total cells (numbers of CD3+ or CC3+ cells divided by numbers of DAPI+ nuclei). Controls were either no T cells (black) or mock T cells (blue). **(E)** IL13Rα2-CAR T cells do not kill PBT308 GOSCs that lack IL13Rα2 expression. Representative confocal images (z-axis projections) of immunostaining for IL13Rα2 (top, white), CD3 (bottom, white) and CC3 (bottom, red) for PBT308 after 7 days of co-culture with either no T cells added, mock T cells, or IL13Rα2-CAR T cells. DAPI (blue) shows nuclei. E:T ratio = 1:1. Scale bar = 50 µm. **(F)** Confirmation of the absence of IL13Rα2 immunostaining of PBT308 day 28 GOSC and its day 0 parental tumor (confocal z-axis projections of representative fields). Scale bar = 50 µm.

We first evaluated CLTX binding in three HGG patient samples. Biotin-conjugated CLTX peptide was added to fixed patient tumors samples and their GOSCs, followed by incubation with fluorophore-conjugated streptavidin and confocal microscope visualization. We observed CLTX binding to 20-90% of total cells in different sub-regions across different parental tumors, with their corresponding GOSCs exhibiting similar binding levels (**Figure 6B**) (image for PBT249 day 28 GOSC in the upper left panel of **Figure 6C**). We suggest that differences between patient samples in percentages of CLTX-binding cells reflect variation in contributions of non-neuroepithelial-derived cells to the total tumor mass.

We then examined whether HGG-directed CAR T cells incorporating a CLTX tumor cell recognition domain can infiltrate GOSCs and kill tumor cells with target specificity. In this study (**Figure 6C**), we challenged living GOSCs with CLTX-CAR T cells, control mock T cells, and only vehicle, for seven days. After seven days, GOSCs were washed, fixed, and stained as whole mounts with markers for T cells (CD3) and apoptosis (cleaved caspase 3; CC3) for IF analysis.

In GOSCs exposed to CLTX-CAR T cells, we observed significantly increased numbers of CD3+ cells to 7-10% of total DAPI+ cells as compared to 1-3 % for vehicle-only controls (**Figure 6D left panel, red and black bars**), indicating infiltration of CLTX-CAR T cells into GOSCs and killing of CLTX-binding tumor cells (observe upper right panel of Figure 6C). At the same time, infiltration of mock T cells varied independently of CLTX-CAR T cells between GOSCs from different patients (**Figure 6D left panel, blue bars**), suggesting patient-related differences in non-targeted T cell retention that will be examined in future studies.

Staining for the apoptosis marker CC3 after CLTX-CAR T cell exposure showed sharply increased apoptosis as compared to GOSCs exposed to mock T cells or vehicle-only controls (**Figure 6D, right panel**), and further that while mock T cells could effectively infiltrate some GOSCs, even if infiltrated they exhibited minimal killing (numbers of CC3+ cells) at levels that could be comparable to vehicle-only controls (note in particular PBT308 for CLTX-CAR T cells). These observations indicate maintained tumor cell-specific killing by CLTX-CAR T cells in GOSCs.

Additionally, we tested IL13Rα2-targeting CAR T cells against GOSCs derived from slices of PBT308. For this PBT line, IF analysis of IL13Rα2 expression revealed that the PBT308 day 28 GOSCs lacked IL13Rα2 expression (**Figure 6E upper left panel, Figure 6F right panel**), as did its parental tumor (**Figure 6F left panel**). Consistent with this absence of IL13Rα2 expression, no apoptotic CC3+ cells were observed after seven-day co-culture with IL13Rα2-targeting CAR T cells (**Figure 6E lower panels**), despite clear infiltration of CD3+ IL13Rα2-CAR T cells into the GOSC.

These results demonstrate long-term stability of target tumor antigen expression (or lack thereof) by GOSCs and maintained specificity of tumor cell target protein recognition by CAR T cells. These observations highlight the utility of GOSCs as an ex vivo platform for evaluating immunotherapeutic agents for brain tumors.

## DISCUSSION

In this study, we established a long-term (at least 28 days) human glioblastoma organotypic slice culture (GOSC) platform that preserves the cellular composition, microenvironment, and architecture of parental tumors. Importantly, tumor vasculature and immune cells, including T cells, tumor-associated macrophages, and microglia, were also maintained for at least 28 days. There were, however, small quantitative increases in populations of some immune cell types at day 28, revealed in transcriptomic expression profiling (bulk RNA-seq) (**Figure 2D**) and also observed in counts of antigen-positive cells in our immunofluorescence analysis (**Figure 5C**). The cause of these increases, which when present were limited to non-tumor cells, is unclear but may relate to unavoidable tissue injury from slicing and subsequent immune cell activation.

Note that while GOSCs were viable beyond day 28, they have not yet been characterized in detail.

Several factors contributed to successful ex vivo culture in our GOSC model. Minimizing disruption of fresh patient samples during vibratome sectioning by using a sapphire blade with carefully adjusted speed and excursion helped preserve tumor spatial relationships, microenvironment including peritumoral gray and white matter, areas of pseudopalisading necrosis, and vascular structures with enclosed red blood cells. Maintaining GOSCs at 4% O_2_, within the physiological range for brain tumors (1–5%) and much lower than atmospheric O_2_ levels (20–21%) typically present in other platforms [31, 32] (**Supplemental Table S2**), conferred stability that we did not see with culture at higher oxygen levels during preliminary optimization experiments (data not shown), consistent with tumor sensitivities to oxygen concentrations [5, 79]. Additionally, our GBS medium incorporates BrainPhys, a medium specifically designed to support in vitro neuron growth and synaptogenesis [80], and containing 2.5 mM D-glucose, close to the concentration in normal human cerebrospinal fluid [36]. In contrast, other investigators [31] [32] (**Supplemental Table S2**) have used DMEM-, DMEM/F12– and Neurobasal-based media with D-glucose concentrations 10–20 times higher than physiological levels. As persistent high glucose levels cause glucotoxicity in cultured cells [81], particularly neurons [82], and is associated with increased risk of neurodegeneration in diabetic patients [83], the lower more physiological glucose concentration in GBS medium may better support long-term GOSC viability. Results of preliminary experiments performed to optimize these culture conditions (not shown) support this suggestion.

Finally, we demonstrate that GOSCs are an ex vivo platform suitable for modeling CAR T cell therapy and immunotherapies in general. First, in co-culture killing assays, CLTX-CAR T cells showed anti-tumor activity paralleling the extent of CLTX binding, which varied between patients while remaining consistent between parental and GOSCs over the 28-day culture period (**Figure 6B**). Thus, CLTX-CAR T cells were highly effective against PBT249 (**Figure 6C**) and PBT308 GOSCs, but less so against PBT309 (**Figure 6D right**) despite infiltrating PBT309 GOSCs (**Figure 6D left**). Similarly, IL13Rα2-CAR T cells showed minimal activity on PBT308 GOSCs lacking IL13Rα2 expression (**Figure 6E**), as was the PBT308 parental tumor (**Figure 6F**). These observations demonstrate the utility of GOSCs for evaluating the specificity and efficacy of HGG-targeted immunotherapies.

A potential limitation of our GOSC platform is the challenge of tumor biobanking for longitudinal, retrospective, and prospective studies. However, recently published protocols for cryopreservation of brain tissue and organoids indicate that this technical difficulty is not insurmountable [84, 85].

In summary, we established a long-term organotypic slice culture platform that preserves the architecture and cellular complexity of parental HGG tumors for at least 28 days. We propose that this model will be a valuable tool for fundamental and translational research, enabling sequential and combinatorial testing of therapies targeting the multiple components of HGGs.

## ACKNOWLEDGMENTS

We extend our deepest gratitude to the patients who donated their tumor tissues to make this work possible.

This work is supported, in part, by grants from the California Institute for Regenerative Medicine (CIRM CLIN2-10248 and DISC2-16638, C.E.B.), the National Institutes of Health (R01 CA254271, C.E.B., B.B.), the Marcus Foundation, the Ivy Foundation. We thank the following City of Hope research cores: Analytical Flow Cytometry, Light Microscopy Digital Imaging, Integrative Genomics, Electron Microscopy, and Biostatistics, which are supported in part by the City of Hope Research Facilities grant P30CA033572 from the National Institutes of Health. The content is solely the responsibility of the authors and does not necessarily represent the official views of the National Institutes of Health. Some figure panels were prepared using BioRender.com.

## AUTHOR CONTRIBUTIONS

Conceptualization: S.W., M.E.B.

Data curation: n/a

Formal analysis: S.W., C.O., H.N.Z, M.D’A., X.W., A.B., Y.H., M.B.

Funding acquisition: C.E.B., M.E.B.

Investigation: S.W.

Methodology: S.W., M.E.B.

Project administration: n/a

Resources: B.B., C.E.B.

Software: n/a

Supervision: C.E.B., M.E.B.

Validation: n/a

Visualization: S.W.

Writing – original draft: S.W., H.N.Z., H.T.K, M.E.B.

Writing – review & editing: S.W., H.N.Z., H.T.K, M.E.B., C.E.B.

## CONFLICT OF INTEREST

C.E.B. and M.E.B. are inventors on patent no. WO2017066481A1 titled “Chimeric antigen receptors containing a chlorotoxin domain”, which is associated with the CLTX-directed CAR T cell, and for which they receive royalty payments. Additionally, patents associated with the IL13Rα2-directed CAR T cell have been licensed by Mustang Bio., Inc., for which C.E.B. receives royalty payments. The other authors declare no competing interests.

Given their role as Editor, Michael Berens had no involvement in the peer-review of this article and had no access to information regarding its peer-review. Full responsibility for the editorial process for this article was delegated to another journal editor.

## ETHICS

The use of de-identified patient tissues was approved by the City of Hope Institutional Review Board (IRB #07074).

## FUNDING

This work is supported, in part, by grants from the California Institute for Regenerative Medicine (CIRM CLIN2-10248 and DISC2-16638, C.E.B.), the National Institute of Health (R01 CA254271, C.E.B., B.B.), the Marcus Foundation, the Ben and Catherine Ivy Foundation. Research cores are supported in part by City of Hope Research Facilities grant P30CA033572 from the National Institutes of Health: Analytical Flow Cytometry, Light Microscopy Digital Imaging, Integrative Genomics, Electron Microscopy, and Biostatistics.

## DATA AVAILABILITY

Data will be made available on reasonable request to the senior authors: Drs. Michael Barish and Christine Brown.

## FIGURE LEGENDS

**Supplemental Figure S1.**
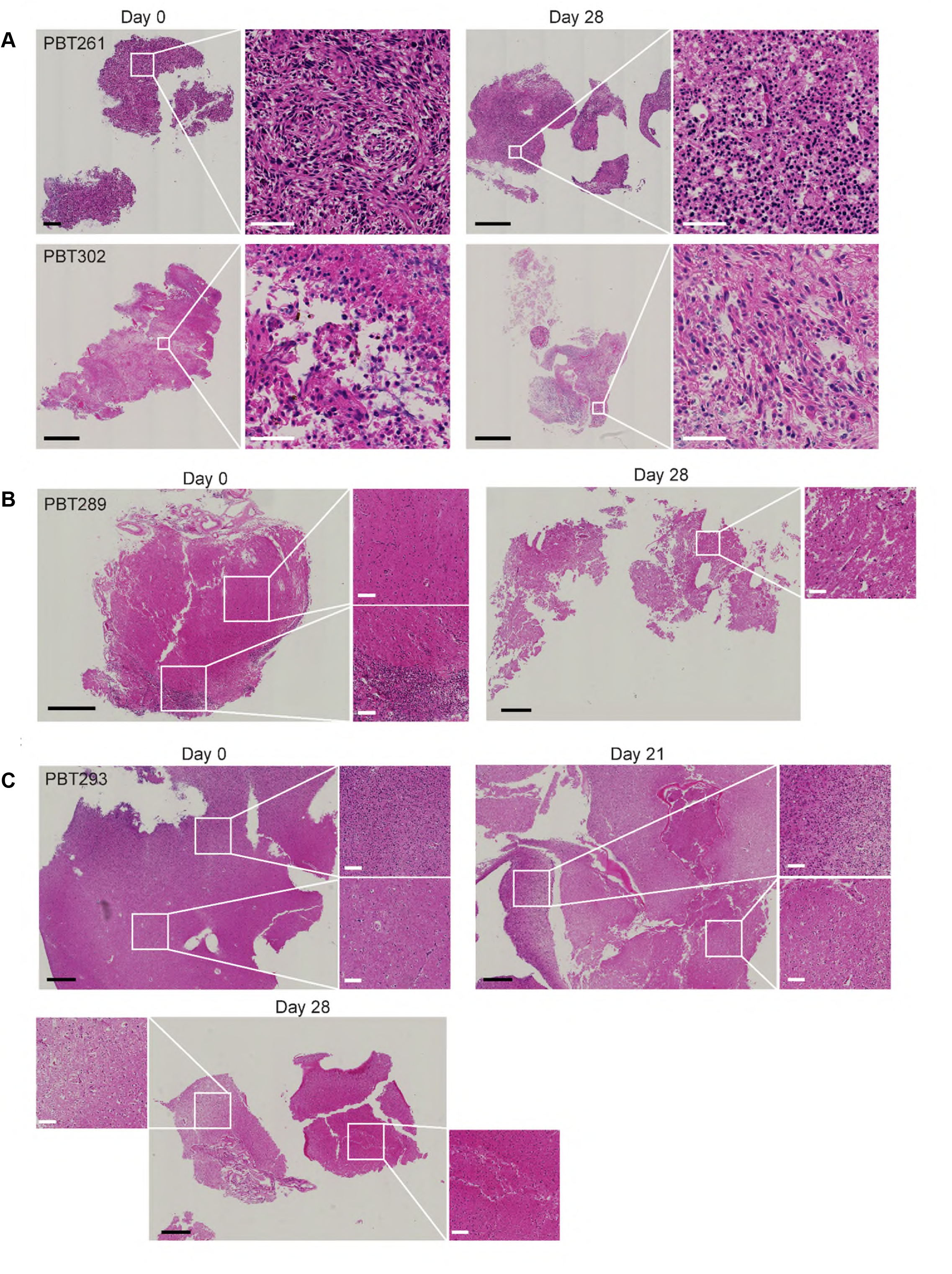
Histopathology features in H&E staining were preserved between day 0 parental tumors and corresponding day 28 GOSCs, related to Figure 1. **(A)** A dense tumor area is preserved in PBT261, and a dispersed tumor is preserved in PBT302. Shown are day 0 parental tumors and corresponding day 28 GOSCs. **(B)** Preservation of non-tumor areas in day 0 parental tumor PBT289 (above dense tumor area at the bottom of the slice), and a non-tumor area in the corresponding day 28 GOSC. **(C)** Preservation of non-tumor areas of day 0 parental tumor PBT293 and corresponding day 21 and day 28 GOSCs. In all panels, black scale bars = 1 mm, and white scale bars (insets) = 50 µm.

**Supplemental Figure S2.**
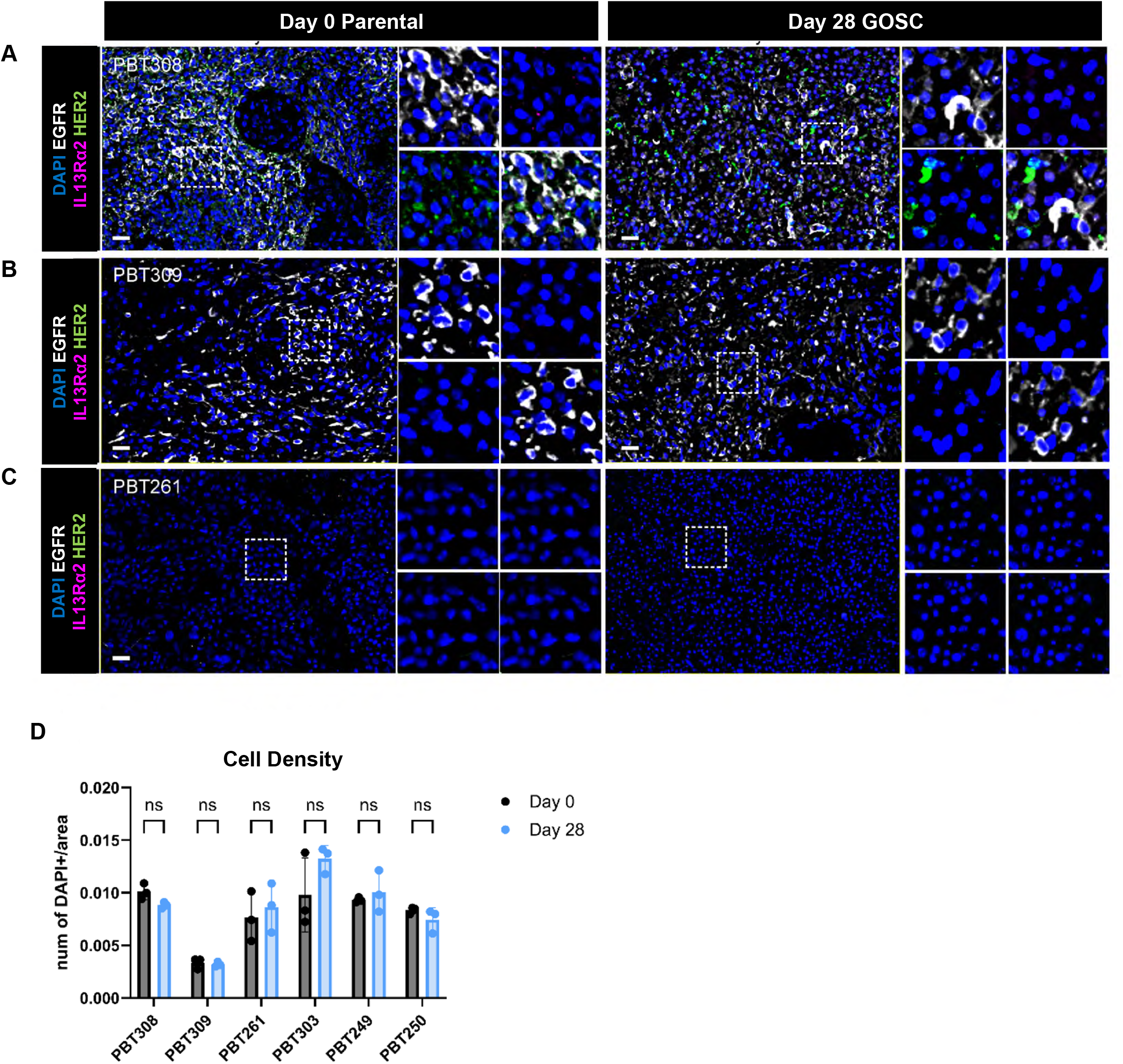
Day 28 GOSCs preserve tumor immunotherapy targets, related to Figure 3A. **(A)** PhenoCycler (CODEX) images of immunostaining in parental day 0 tumors PBT308, PBT309, and PBT261 and corresponding day 28 GOSCs for three tumor immunotherapy targets: HER2 (green), IL13Rα2 (magenta), and EGFR (white). DAPI (blue), nuclei. Scale bar, 50 µm. **(B)** No statically significant differences were seen in quantifications of cell densities between six pairs of day 0 parental tumor samples and day 28 GOSCs. Data (mean ± SD) are presented as number of DAPI+ cells divided by the area of the tissue. **(C)** No statically significant differences were seen in quantifications of EGFR, IL13Rα2, and HER2 expression between day 0 parental tumor samples and day 28 GOSCs. Data (mean ± SD) are presented as area of positive expression (pixels, as µm^2^) divided by the total area (µm^2^).

**Supplemental Figure S3.**
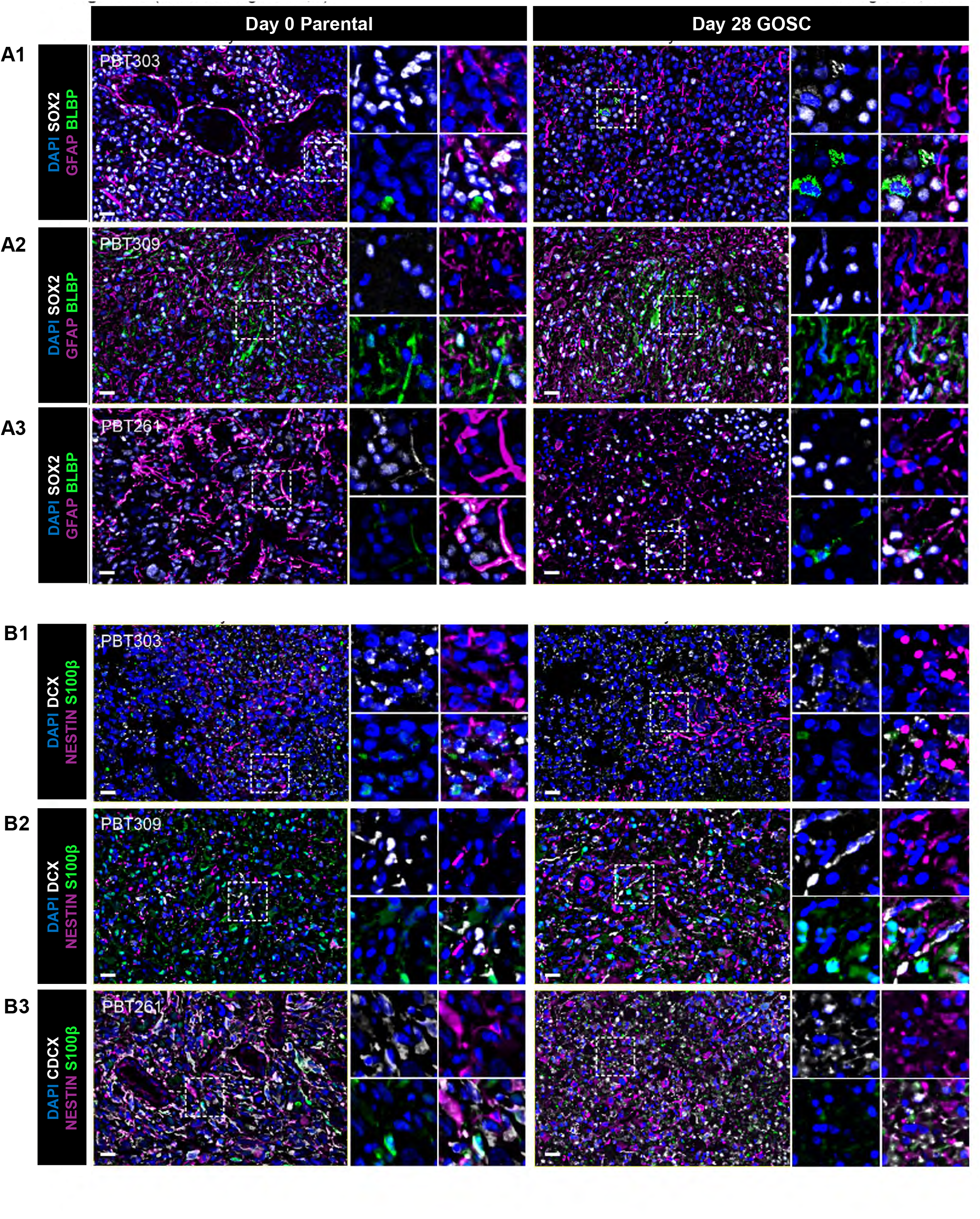
Day 28 GOSCs preserve glioma stem/progenitor cells and neuronal lineage cells from day 0 parental tumors, related to Figure 3B and 3C. **(A)** PhenoCycler (CODEX) images of nuclei (DAPI, blue) and immunostaining for early developmental markers brain lipid binding protein (BLBP, green), GFAP (magenta), and SOX2 (white), on day 0 parental tumors (PBT303, PBT309, PBT261) and corresponding day 28 GOSCs. Scale bar = 20 µm. **(B)** PhenoCycler (CODEX) images of nuclei (DAPI, blue) and immunostaining for maturing developmental markers S100β (green), NESTIN (magenta), and doublecortin (DCX, white), and DAPI (blue) on day 0 parental tumors (PBT303, PBT309, PBT261) and corresponding day 28 GOSCs. Scale bar = 20 µm.

**Supplemental Figure S4.**
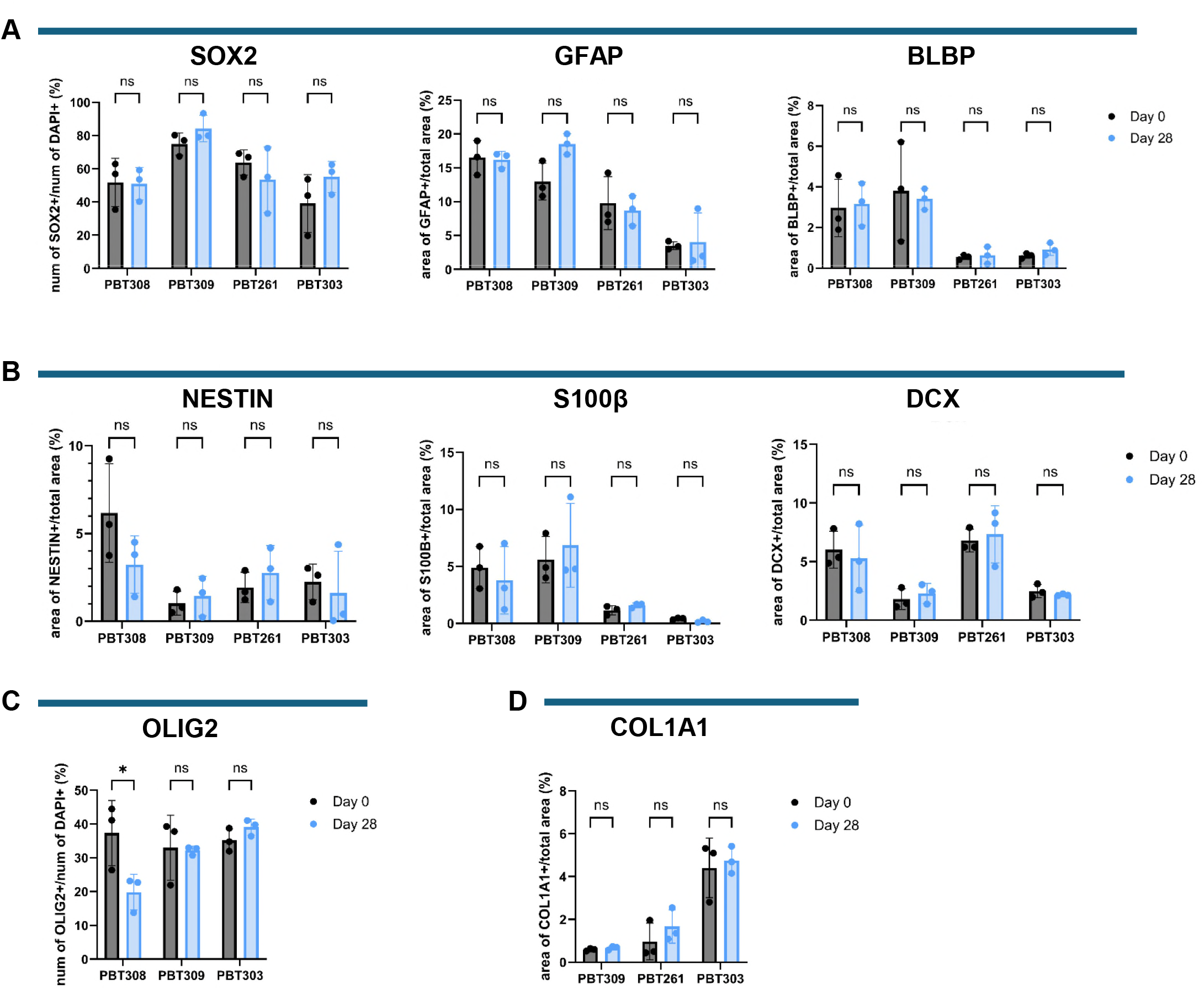
Stable quantified expression of multiple marker expression on day 0 parental tumors and day 28 GOSCs, related to Figure 3B-3E and Supplemental Figure S4. **(A-D)** No statistically significant differences were seen between quantifications (mean ± SD) of expression on day 0 parental tumors and corresponding day 28 GOSCs for **(A)** early developmental markers SOX2, GFAP, and BLBP, **(B)** NESTIN, S100β, and DCX, **(C)** oligodendrocyte lineage marker OLIG2, and **(D)** extracellular matrix component collagen 1 A1 (COL1A1). Data are presented as number of positive cells divided by number of DAPI+ cells for nucleus-localized markers, and areas of positive expression (pixels, presented as µm^2^) divided by the total area (µm^2^) for markers expressed in cytoplasm.

**Supplemental Figure S5.**
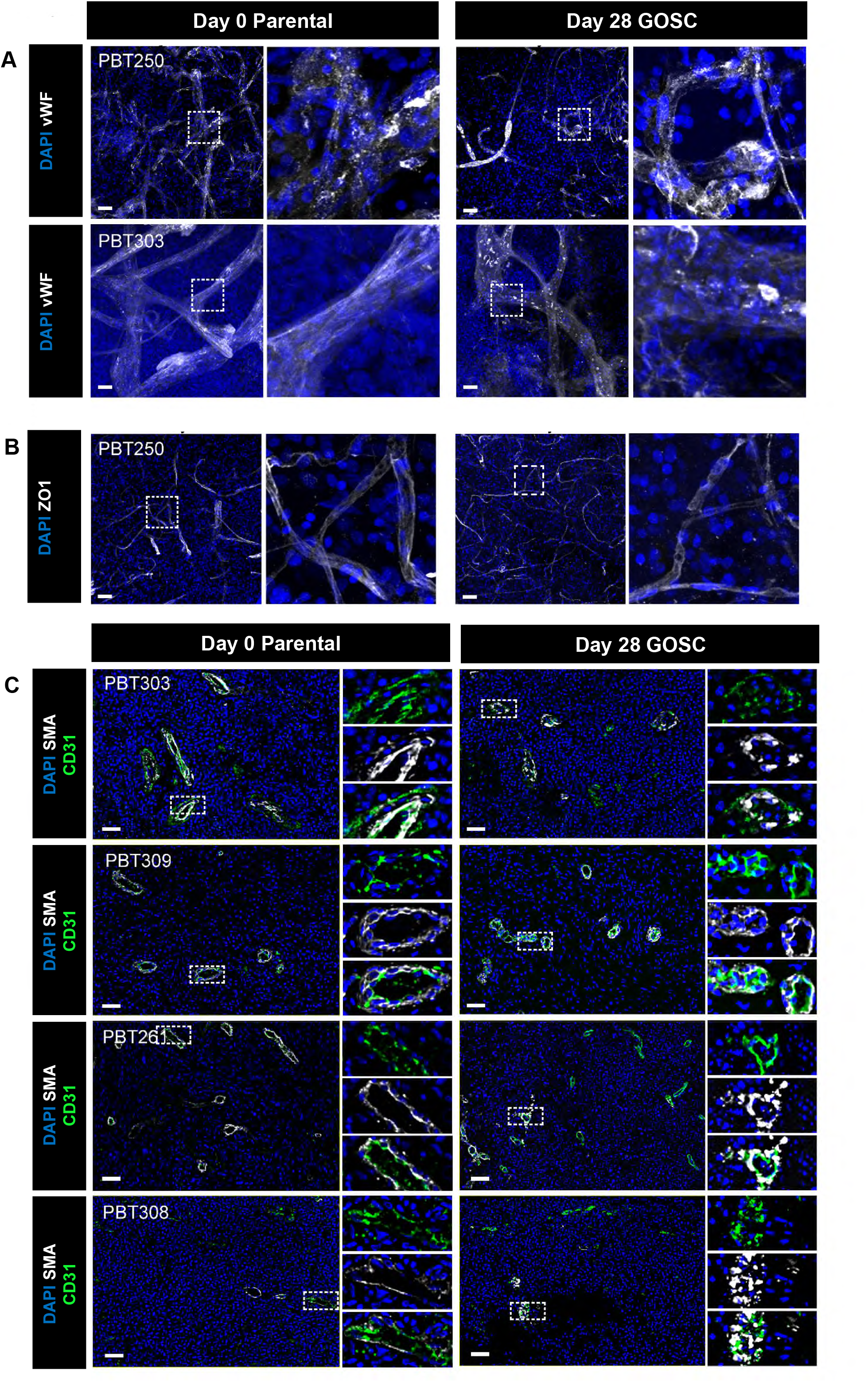
Day 28 GOSCs preserve the tumor vasculature of day 0 parental tumors, related to Figure 4. **(A)** Confocal images (z-axis projection) of immunostaining for von Willebrand factor (vWF, white) in day 0 parental tumors and corresponding day 28 GOSCs. DAPI, blue. Scale bar, 50 µm. **(B)** Confocal images (z-axis projection) of immunostaining for ZO1 (white) on day 0 parental tumor PBT250 and corresponding day 28 GOSC. DAPI, blue. Scale bar, 50 µm. **(C)** PhenoCycler (CODEX) images for CD31 (green) and SMA (white) on four day 0 parental tumors (PBT303, PBT309, PBT261, PBT308) and corresponding day 28 GOSCs. DAPI, blue. Scale bar = 50 µm.

**Supplemental Figure S6.**
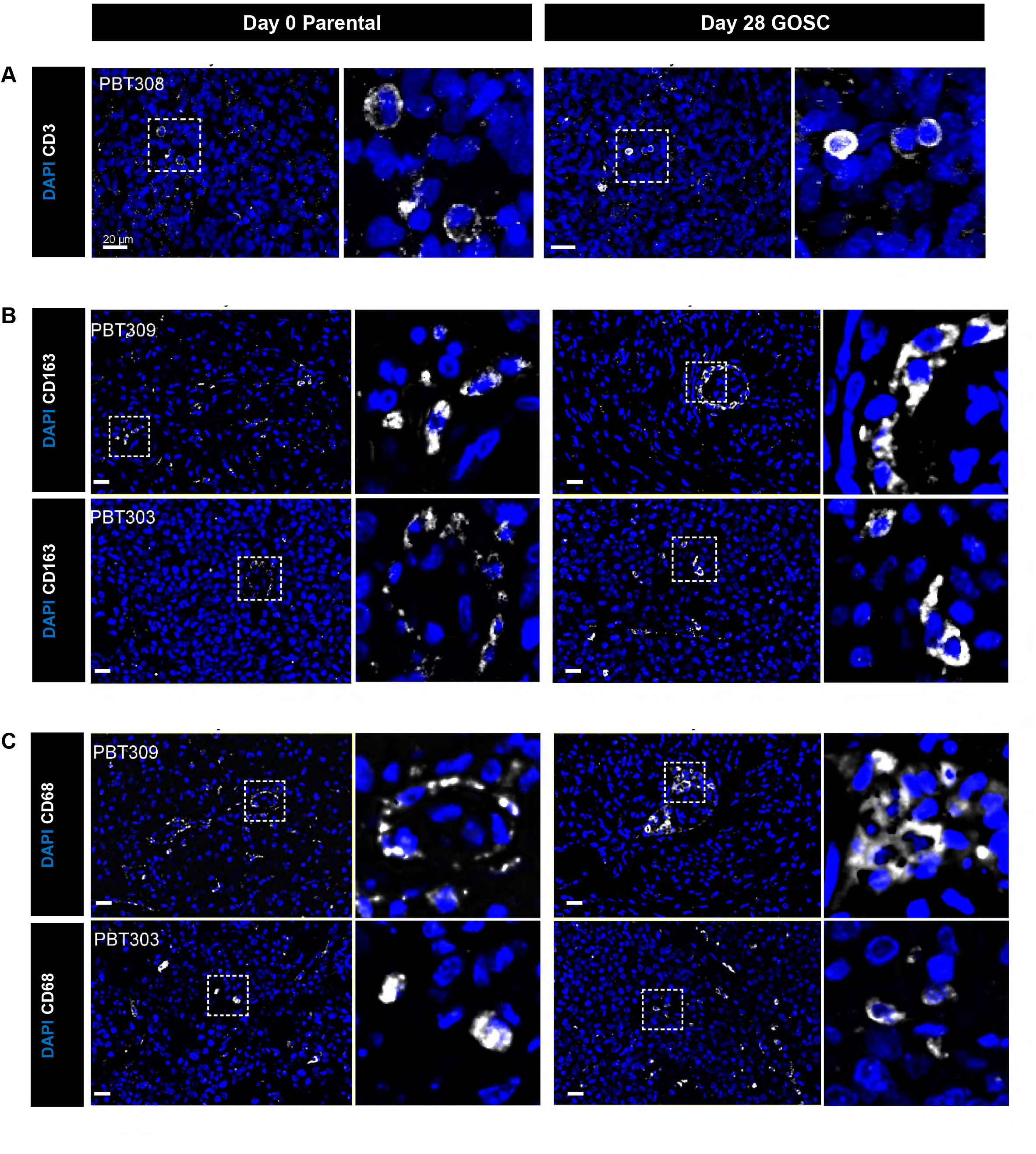
Day 28 GOSCs preserve immune cells from parental day 0 tumors, related to Figure 5. **(A)** Confocal images (z-axis projection) of immunostaining for CD3 (white) on day 0 parental tumor PBT308 and corresponding day 28 GOSC. DAPI, blue. Scale bar = 20 µm. **(B)** PhenoCycler (CODEX) images of immunostaining for CD163 (white) on day 0 parental tumors PBT309 and PBT303, and corresponding day 28 GOSCs. DAPI, blue. Scale bar = 20 µm. **(C)** PhenoCycler (CODEX) images of immunostaining for CD68 (white) on day 0 parental tumors PBT309 and PBT303, and corresponding day 28 GOSCs. DAPI, blue. Scale bar = 20 µm.

**Supplemental Figures S7 and S8.**
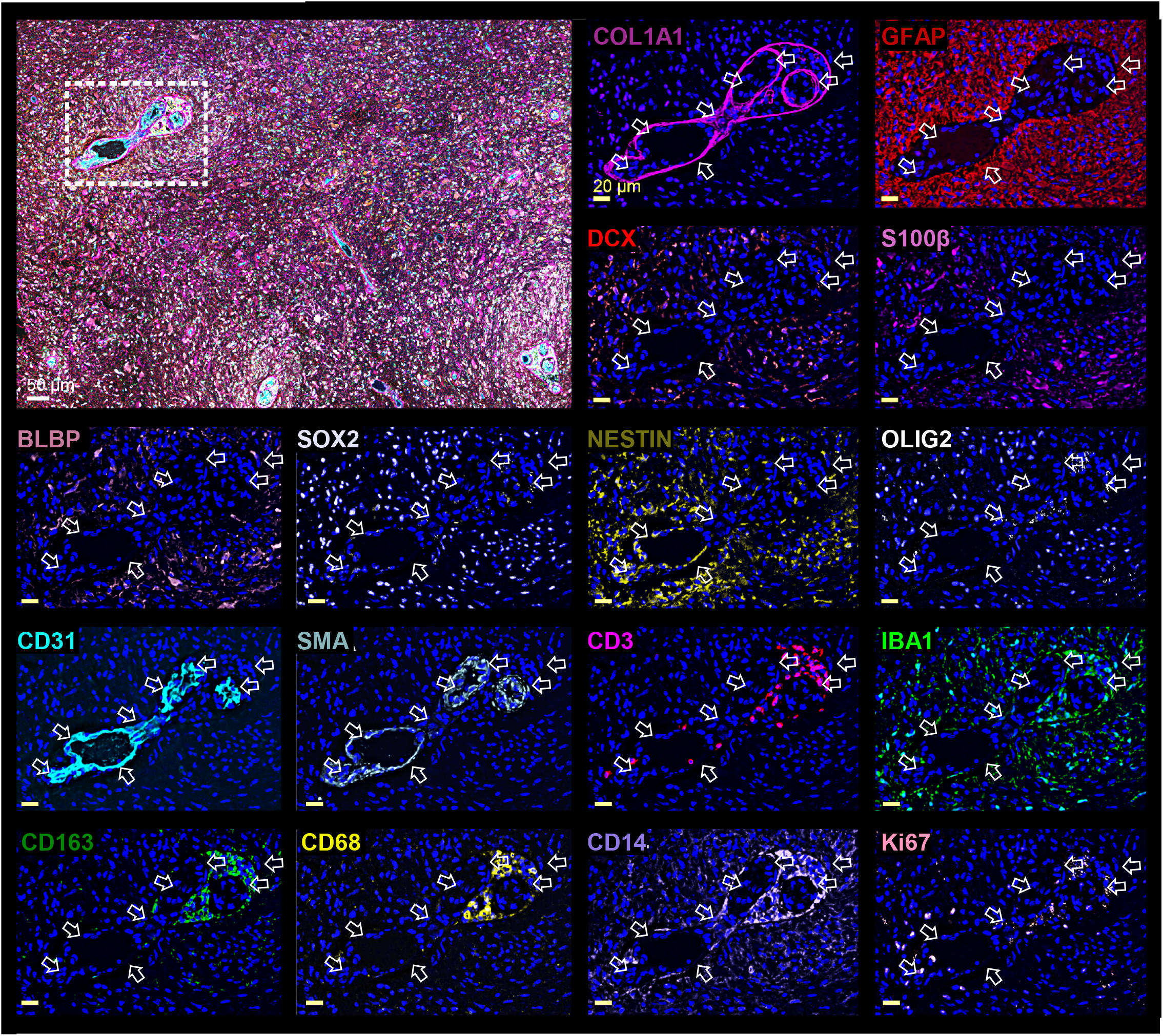
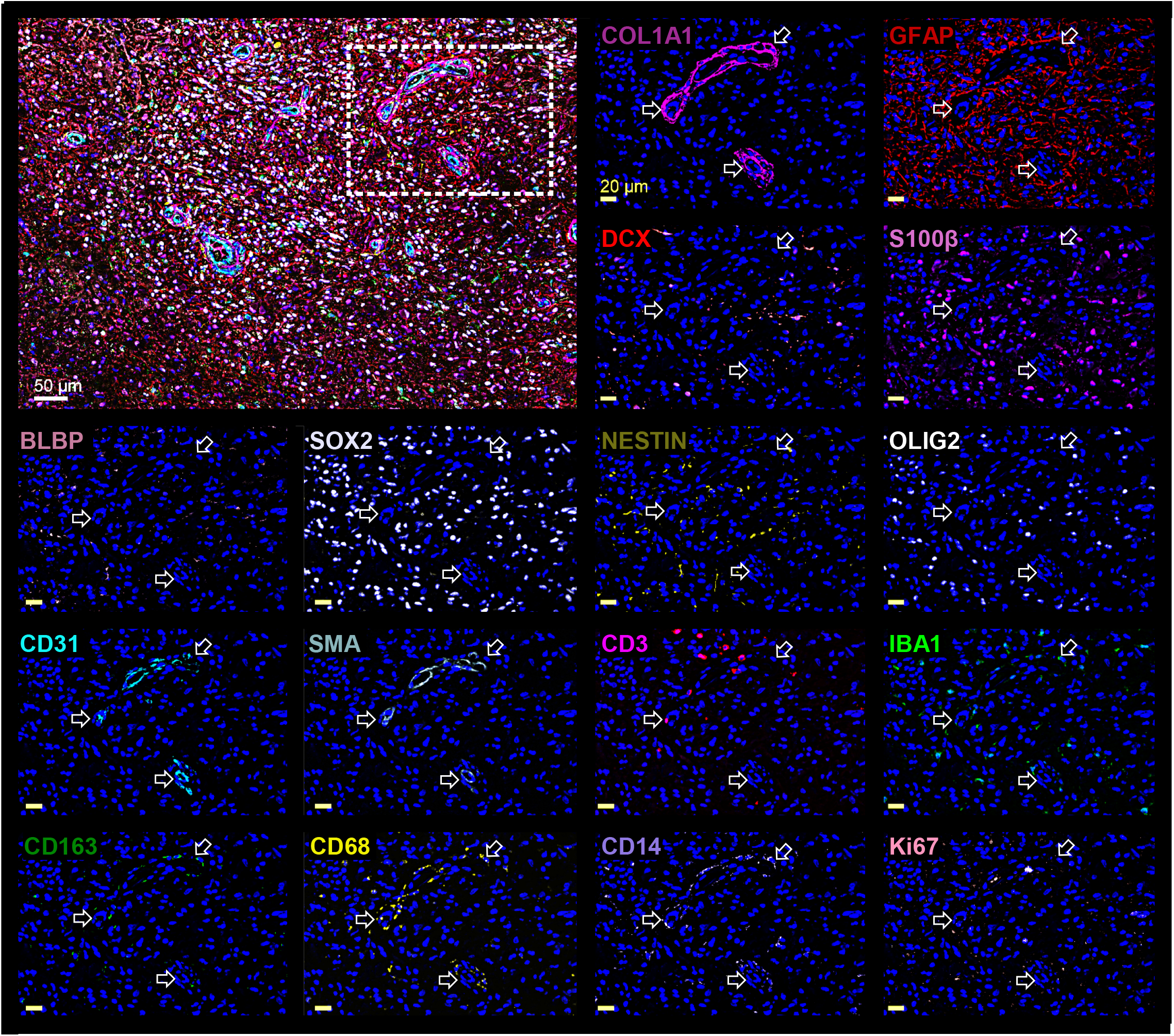
Phenocycler (CODEX) images of day 28 PBT309 GOSC (S7) and of corresponding day 0 PBT309 parental tumor (S8) showing preserved spatial distributions of tumor and microenvironment cells in relation to positions of vasculature within the tumor mass. PhenoCycler (CODEX) multiplexed immunostaining of (**S7**) PBT309 day 28 GOSC, and (**S8**) day 0 PBT309 parental tumor. In each figure, top left is the merged image of 16 markers. Subsequent images display the area within the inset: DAPI (blue) staining of nuclei, and single markers (from left to right, top to bottom): COL1A1 (deep purple), GFAP (dark red), DCX (peach), S100β (magenta), BLBP (pink), SOX2 (leaf green), NESTIN (deep green), OLIG2 (white), CD31 (teal), SMA (blue-green), CD3 (red), IBA1 (light green), CD163 (green), CD68 (yellow), CD14 (purple), and Ki67 (light pink). White scale bars = 50 µm. Yellow scale bars = 20 µm. In **S7** and **S8**, arrows are placed at various positions around CD31 immunofluorescent structures marking vascular endothelial cells. Positions of DAPI+ nuclei relative to these arrows were noted and used as fiduciary points for placement of arrows in the panels from the same Phenocycler (CODEX) image. In the GOSC tumor (**S7**), CD31+ structures marked a large vessel in the lower left and structures with smaller lumens and less compactly arranged endothelial cells in the upper right. SMA+ structures were closely associated with both large and smaller vessels. Circulating immune cells (CD3+, CD163+, CD68+, CD14+) were preferentially clustered in the area of less compact CD31+ and SMA+ vasculature. IBA1+ microglia and macrophages were also associated with less compact vasculature as well as distributed throughout the tumor tissue. A tight band of extracellular matrix component COL1A1 was located outside (5-10 um) vascular structures. Cell types comprising progenitor and NSC-derived lineages (SOX2+, BLBP+, DCX+, OLIG2+, S100β+) were found in tumor tissue. NESTIN+ cells were both around the larger vasculature structure and in clusters within the tumor. GFAP+ astrocytes and tumor cells were excluded from vasculature-rich areas. Cells in the parental tumor (**S8**) were organized similarly although in some cases were not as abundant in these images. CD31+ and SMA+ structures marked vasculature and COL1A1+ extracellular matrix was close to but not overlapping vascular components. Immune cells were found associated with vasculature and progenitor/NSC cells arrayed throughout the tumor.

---

**SUPPLEMENTAL TABLE 1:**
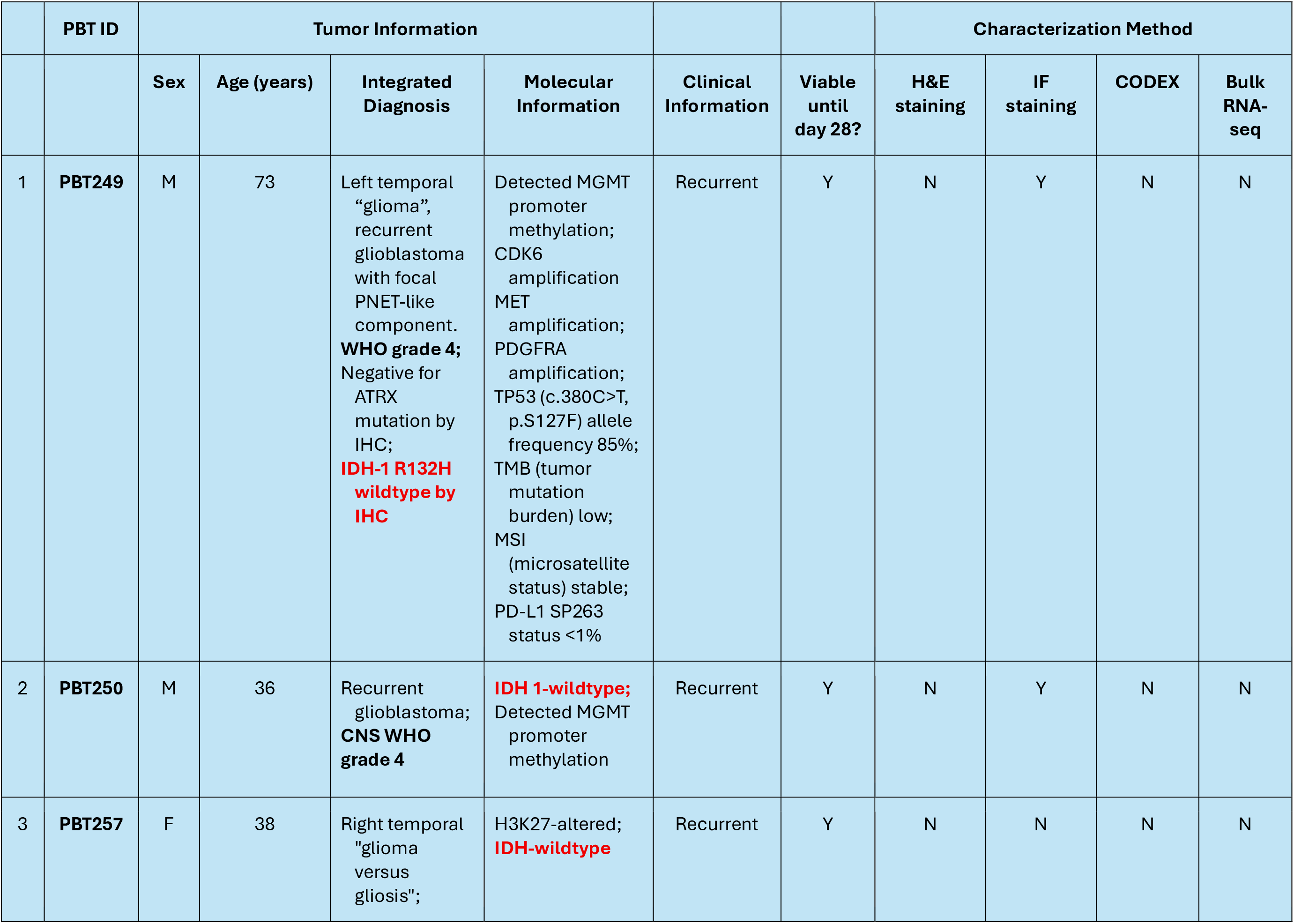
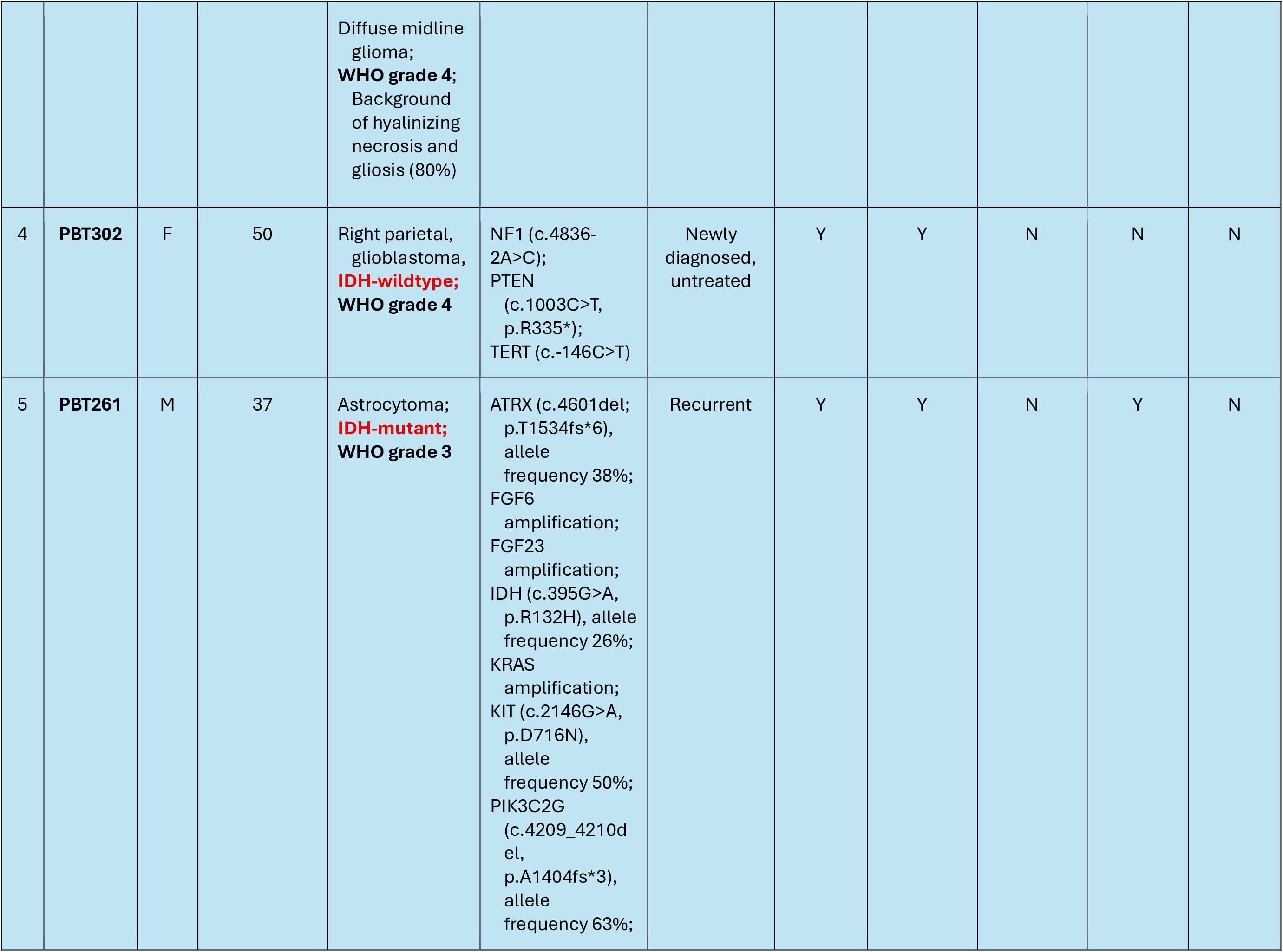
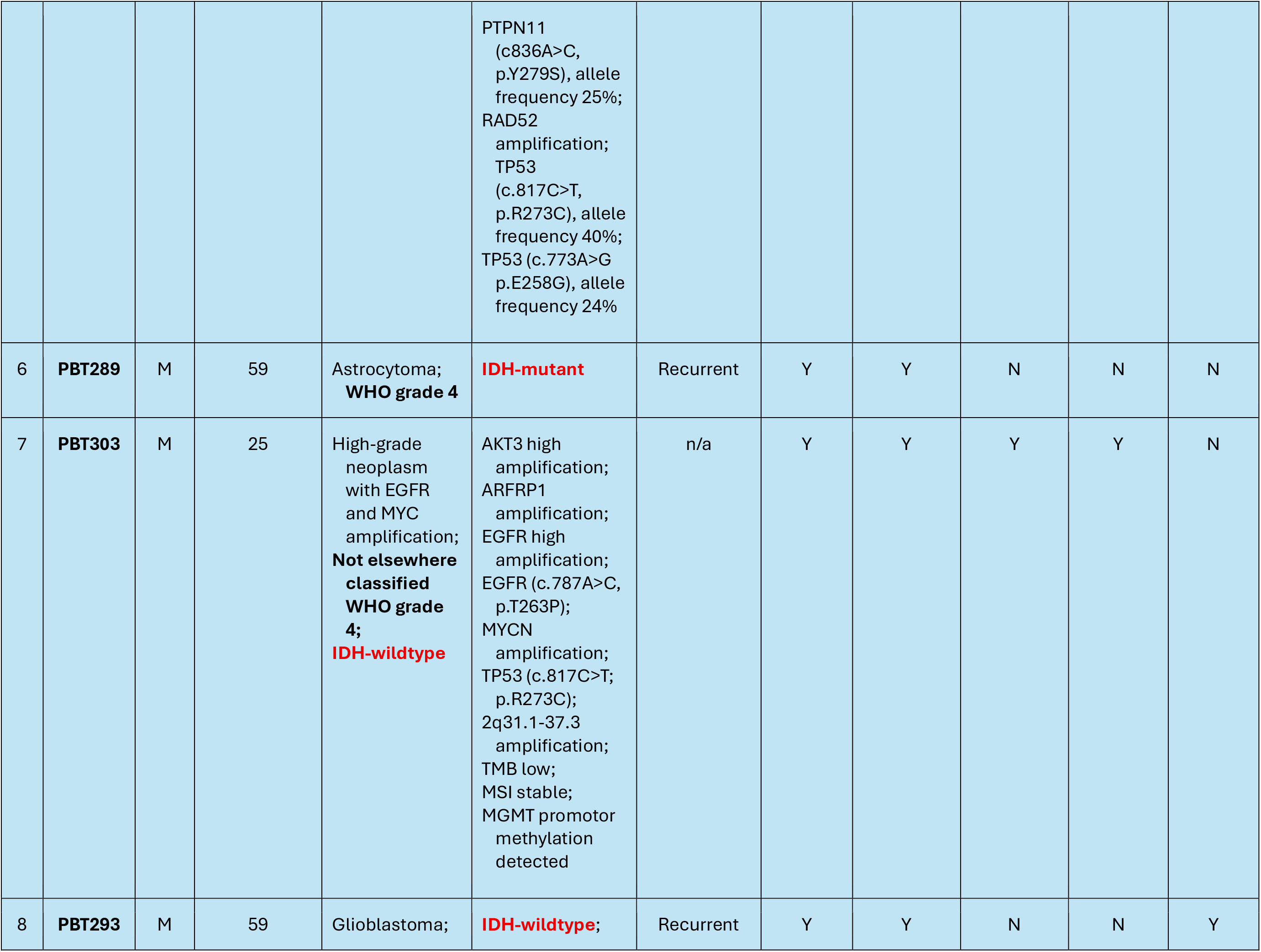
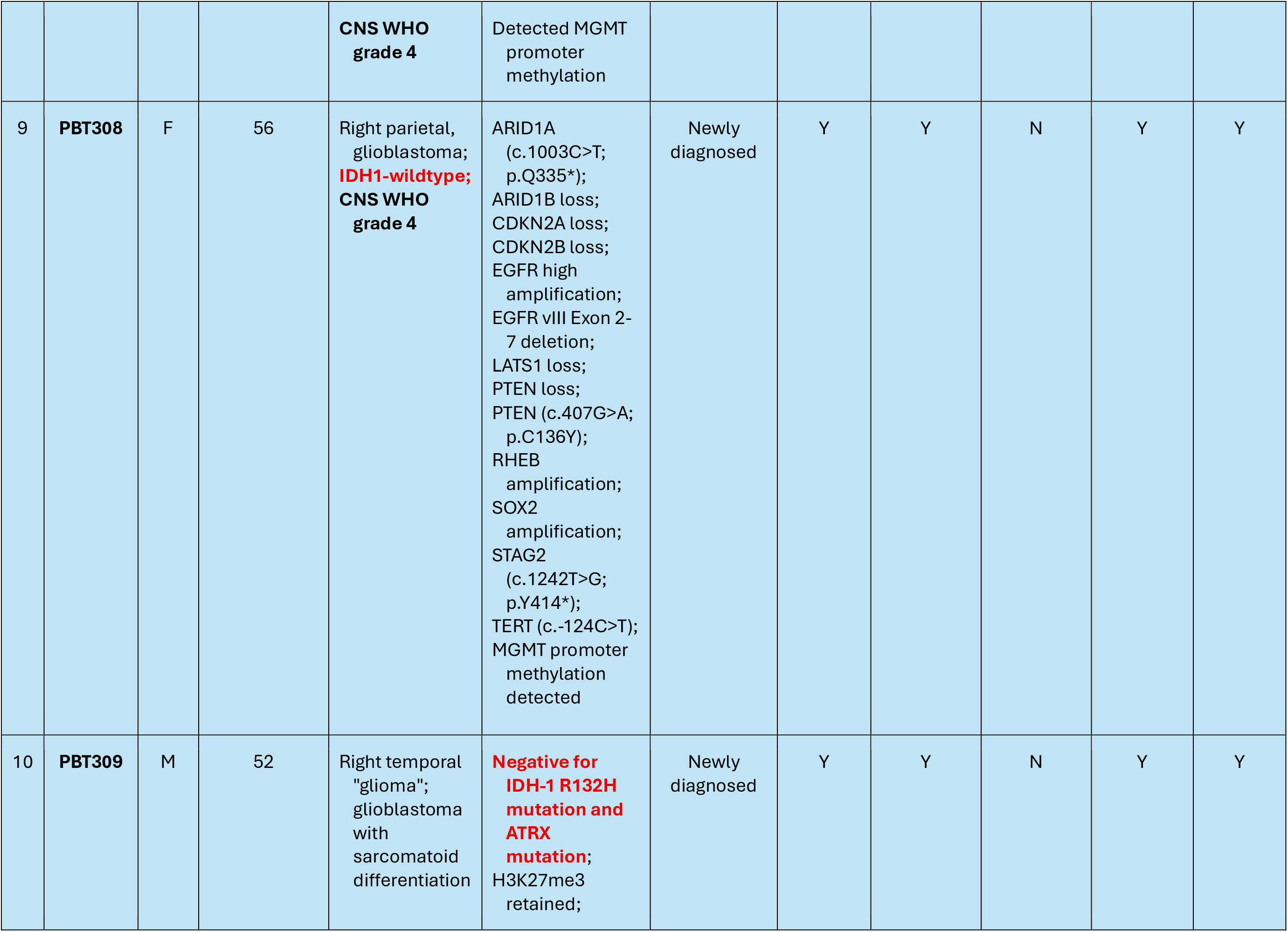
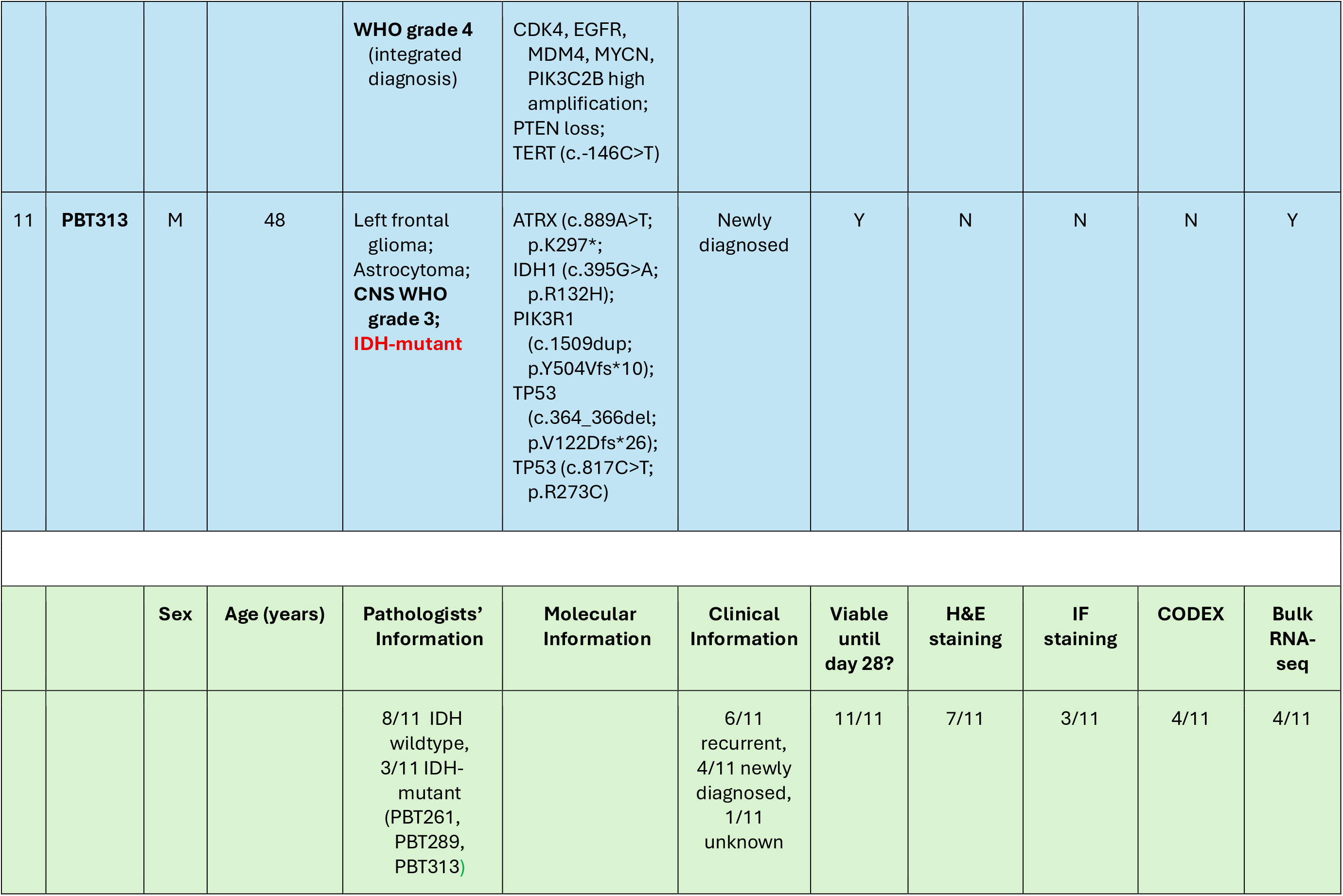
INFORMATION ON TUMOR SAMPLES.

|  | PBT ID | Tumor Information |  |  |  |  |  | Characterization Method |  |  |  |
| --- | --- | --- | --- | --- | --- | --- | --- | --- | --- | --- | --- |
|  |  | Sex | Age (years) | Integrated Diagnosis | Molecular Information | Clinical Information | Viable until day 28? | H&E staining | IF staining | CODEX | Bulk RNA-seq |
| 1 | PBT249 | M | 73 | Left temporal "glioma", recurrent glioblastoma with focal PNET-like component. **WHO grade 4;** Negative for ATRX mutation by IHC; **IDH-1 R132H wildtype by IHC** | Detected MGMT promoter methylation; CDK6 amplification; MET amplification; PDGFRA amplification; TP53 (c.380C>T, p.S127F) allele frequency 85%; TMB (tumor mutation burden) low; MSI (microsatellite status) stable; PD-L1 SP263 status <1% | Recurrent | Y | N | Y | N | N |
| 2 | PBT250 | M | 36 | Recurrent glioblastoma; **CNS WHO grade 4** | **IDH 1-wildtype;** Detected MGMT promoter methylation | Recurrent | Y | N | Y | N | N |
| 3 | PBT257 | F | 38 | Right temporal "glioma versus gliosis"; | H3K27-altered; **IDH-wildtype** | Recurrent | Y | N | N | N | N |
|  |  |  |  | Diffuse midline glioma;<br>**WHO grade 4**;<br>Background of hyalinizing necrosis and gliosis (80%) |  |  |  |  |  |  |  |
| 4 | **PBT302** | F | 50 | Right parietal, glioblastoma, **IDH-wildtype**;<br>**WHO grade 4** | NF1 (c.4836-2A>C); PTEN (c.1003C>T, p.R335*); TERT (c.-146C>T) | Newly diagnosed, untreated | Y | Y | N | N | N |
| 5 | **PBT261** | M | 37 | Astrocytoma; **IDH-mutant**;<br>**WHO grade 3** | ATRX (c.4601del; p.T1534fs*6), allele frequency 38%; FGF6 amplification; FGF23 amplification; IDH (c.395G>A, p.R132H), allele frequency 26%; KRAS amplification; KIT (c.2146G>A, p.D716N), allele frequency 50%; PIK3C2G (c.4209_4210del, p.A1404fs*3), allele frequency 63%; | Recurrent | Y | Y | N | Y | N |
|  |  |  |  |  | PTPN11 (c836A>C, p.Y279S), allele frequency 25%;<br>RAD52 amplification;<br>TP53 (c.817C>T, p.R273C), allele frequency 40%;<br>TP53 (c.773A>G p.E258G), allele frequency 24% |  |  |  |  |  |  |
| 6 | PBT289 | M | 59 | Astrocytoma;<br>**WHO grade 4** | **IDH-mutant** | Recurrent | Y | Y | N | N | N |
| 7 | PBT303 | M | 25 | High-grade neoplasm with EGFR and MYC amplification;<br>**Not elsewhere classified WHO grade 4; IDH-wildtype** | AKT3 high amplification;<br>ARFRP1 amplification;<br>EGFR high amplification;<br>EGFR (c.787A>C, p.T263P);<br>MYCN amplification;<br>TP53 (c.817C>T; p.R273C);<br>2q31.1-37.3 amplification;<br>TMB low;<br>MSI stable;<br>MGMT promotor methylation detected | n/a | Y | Y | Y | Y | N |
| 8 | PBT293 | M | 59 | Glioblastoma; | **IDH-wildtype;** | Recurrent | Y | Y | N | N | Y |

|  |  |  |  | **CNS WHO**<br>**grade 4** | Detected MGMT<br>promoter<br>methylation |  |  |  |  |  |  |
| --- | --- | --- | --- | --- | --- | --- | --- | --- | --- | --- | --- |
| 9 | **PBT308** | F | 56 | Right parietal,<br>glioblastoma;<br>**IDH1-wildtype;**<br>**CNS WHO**<br>**grade 4** | ARID1A<br>(c.1003C>T;<br>p.Q335*);<br>ARID1B loss;<br>CDKN2A loss;<br>CDKN2B loss;<br>EGFR high<br>amplification;<br>EGFR vIII Exon 2-7 deletion;<br>LATS1 loss;<br>PTEN loss;<br>PTEN (c.407G>A;<br>p.C136Y);<br>RHEB<br>amplification;<br>SOX2<br>amplification;<br>STAG2<br>(c.1242T>G;<br>p.Y414*);<br>TERT (c.-124C>T);<br>MGMT promoter<br>methylation<br>detected | Newly<br>diagnosed | Y | Y | N | Y | Y |
| 10 | **PBT309** | M | 52 | Right temporal<br>"glioma";<br>glioblastoma<br>with<br>sarcomatoid<br>differentiation | **Negative for**<br>**IDH-1 R132H**<br>**mutation and**<br>**ATRX**<br>**mutation;**<br>H3K27me3<br>retained; | Newly<br>diagnosed | Y | Y | N | Y | Y |
|  |  |  |  | **WHO grade 4**<br>(integrated diagnosis) | CDK4, EGFR, MDM4, MYCN, PIK3C2B high amplification; PTEN loss; TERT (c.-146C>T) |  |  |  |  |  |  |
| 11 | **PBT313** | M | 48 | Left frontal glioma; Astrocytoma; **CNS WHO grade 3; IDH-mutant** | ATRX (c.889A>T; p.K297*; IDH1 (c.395G>A; p.R132H); PIK3R1 (c.1509dup; p.Y504Vfs*10); TP53 (c.364_366del; p.V122Dfs*26); TP53 (c.817C>T; p.R273C) | Newly diagnosed | Y | N | N | N | Y |
|  |  | **Sex** | **Age (years)** | **Pathologists' Information** | **Molecular Information** | **Clinical Information** | **Viable until day 28?** | **H&E staining** | **IF staining** | **CODEX** | **Bulk RNA-seq** |
|  |  |  |  | 8/11 IDH wildtype, 3/11 IDH-mutant (PBT261, PBT289, PBT313) |  | 6/11 recurrent, 4/11 newly diagnosed, 1/11 unknown | 11/11 | 7/11 | 3/11 | 4/11 | 4/11 |

**SUPPLEMENTAL TABLE 2:**
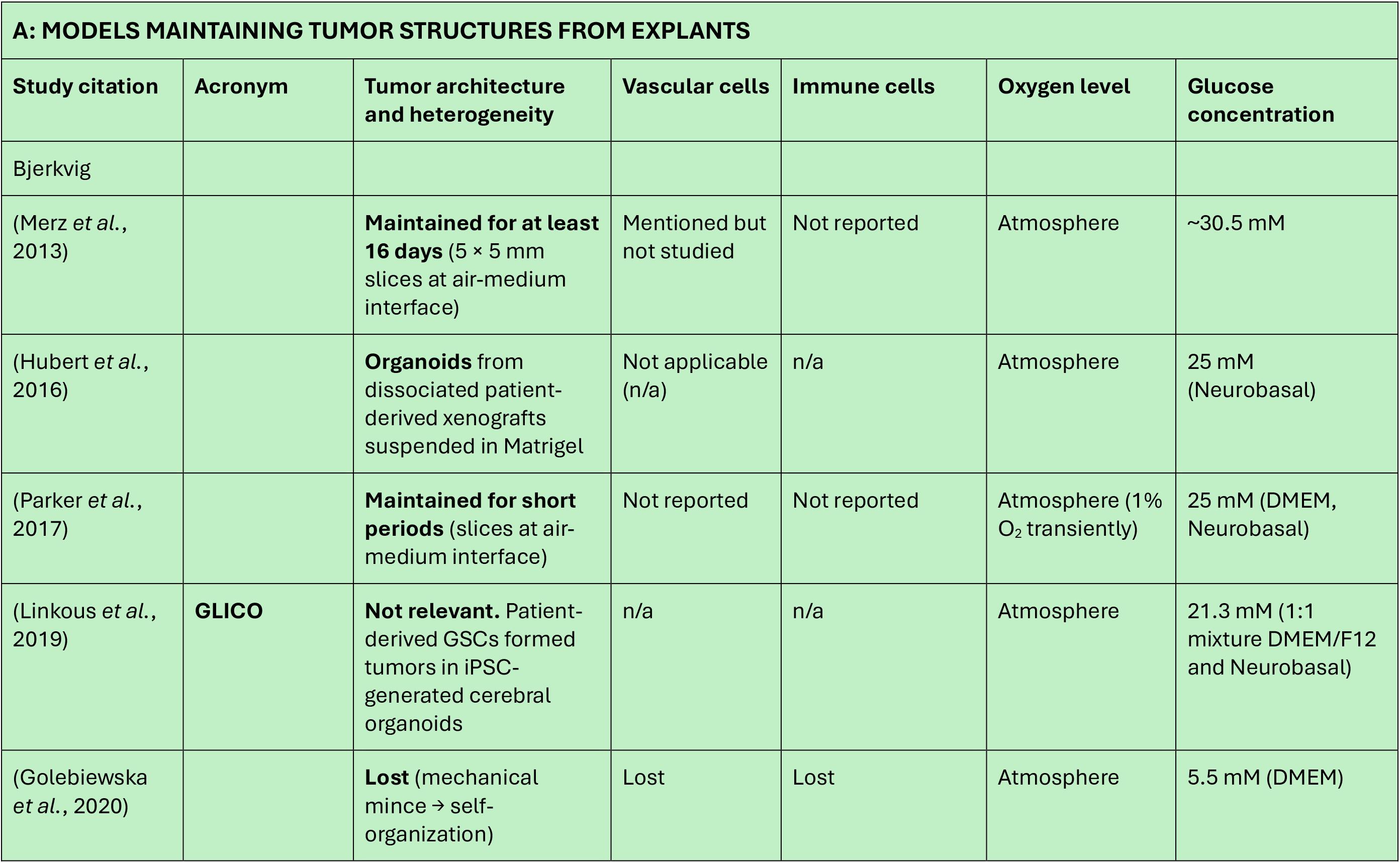
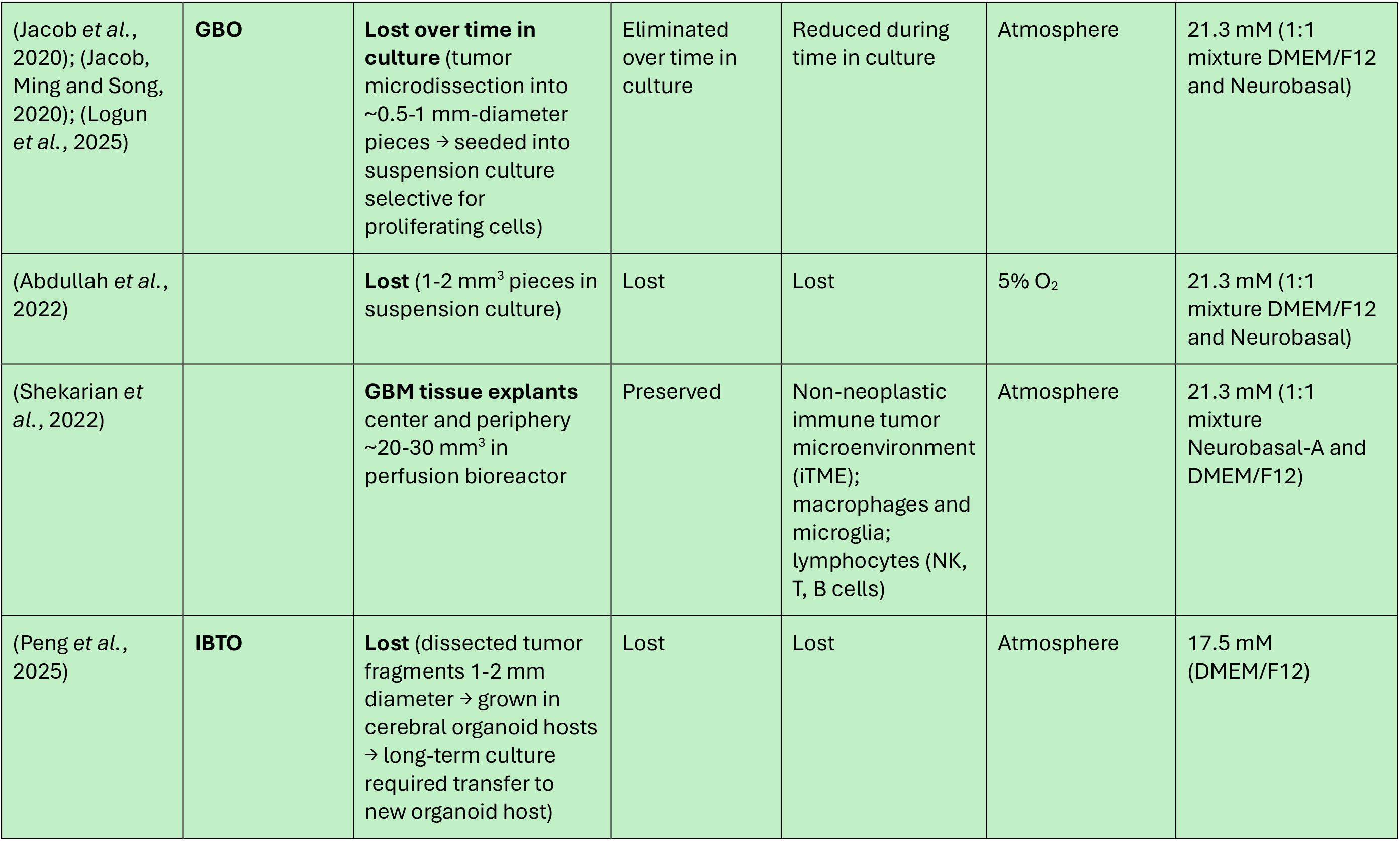
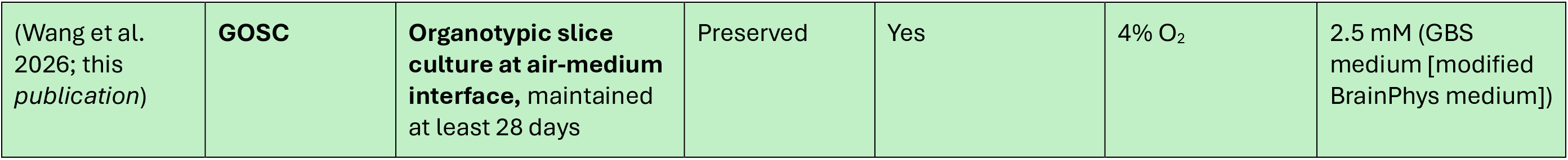
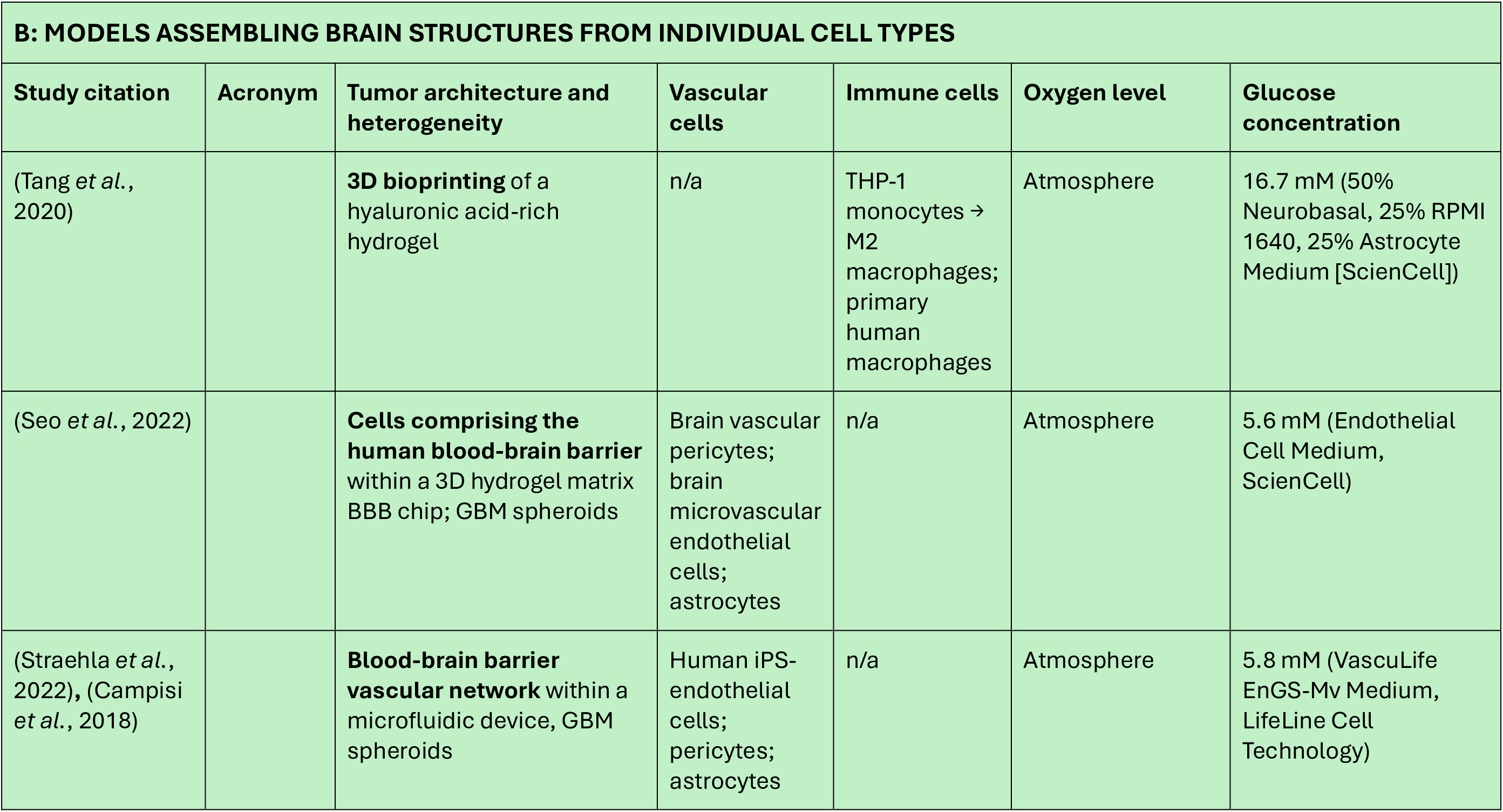
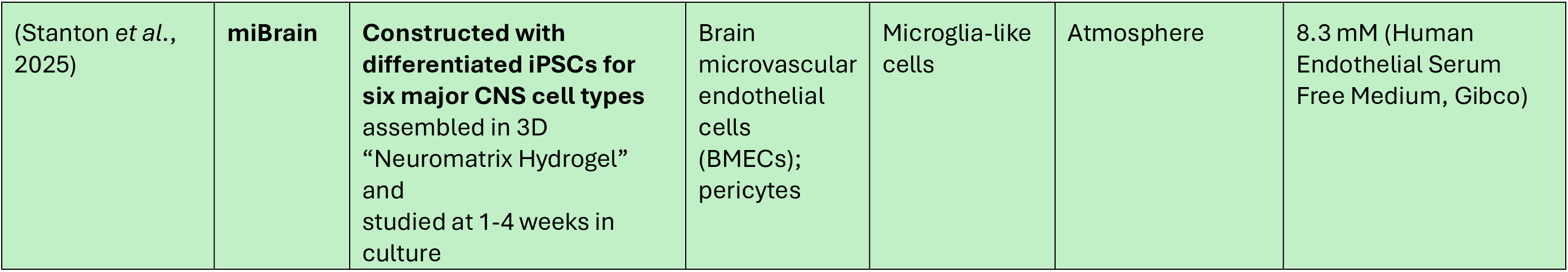
OTHER EX VIVO BRAIN AND HGG MODELS.

| A: MODELS MAINTAINING TUMOR STRUCTURES FROM EXPLANTS |  |  |  |  |  |  |
| --- | --- | --- | --- | --- | --- | --- |
| Study citation | Acronym | Tumor architecture and heterogeneity | Vascular cells | Immune cells | Oxygen level | Glucose concentration |
| Bjerkvig |  |  |  |  |  |  |
| (Merz *et al.*, 2013) |  | **Maintained for at least 16 days** (5 × 5 mm slices at air-medium interface) | Mentioned but not studied | Not reported | Atmosphere | ~30.5 mM |
| (Hubert *et al.*, 2016) |  | **Organoids** from dissociated patient-derived xenografts suspended in Matrigel | Not applicable (n/a) | n/a | Atmosphere | 25 mM (Neurobasal) |
| (Parker *et al.*, 2017) |  | **Maintained for short periods** (slices at air-medium interface) | Not reported | Not reported | Atmosphere (1% O<sub>2</sub> transiently) | 25 mM (DMEM, Neurobasal) |
| (Linkous *et al.*, 2019) | **GLICO** | **Not relevant.** Patient-derived GSCs formed tumors in iPSC-generated cerebral organoids | n/a | n/a | Atmosphere | 21.3 mM (1:1 mixture DMEM/F12 and Neurobasal) |
| (Golebiewska *et al.*, 2020) |  | **Lost** (mechanical mince → self-organization) | Lost | Lost | Atmosphere | 5.5 mM (DMEM) |
| (Jacob *et al.*, 2020); (Jacob, Ming and Song, 2020); (Logun *et al.*, 2025) | **GBO** | **Lost over time in culture** (tumor microdissection into ~0.5-1 mm-diameter pieces → seeded into suspension culture selective for proliferating cells) | Eliminated over time in culture | Reduced during time in culture | Atmosphere | 21.3 mM (1:1 mixture DMEM/F12 and Neurobasal) |
| (Abdullah *et al.*, 2022) |  | **Lost** (1-2 mm<sup>3</sup> pieces in suspension culture) | Lost | Lost | 5% O<sub>2</sub> | 21.3 mM (1:1 mixture DMEM/F12 and Neurobasal) |
| (Shekarian *et al.*, 2022) |  | **GBM tissue explants** center and periphery ~20-30 mm<sup>3</sup> in perfusion bioreactor | Preserved | Non-neoplastic immune tumor microenvironment (iTME); macrophages and microglia; lymphocytes (NK, T, B cells) | Atmosphere | 21.3 mM (1:1 mixture Neurobasal-A and DMEM/F12) |
| (Peng *et al.*, 2025) | **IBTO** | **Lost** (dissected tumor fragments 1-2 mm diameter → grown in cerebral organoid hosts → long-term culture required transfer to new organoid host) | Lost | Lost | Atmosphere | 17.5 mM (DMEM/F12) |
| (Wang et al. 2026; this *publication*) | **GOSC** | **Organotypic slice culture at air-medium interface**, maintained at least 28 days | Preserved | Yes | 4% O<sub>2</sub> | 2.5 mM (GBS medium [modified BrainPhys medium]) |

| **B: MODELS ASSEMBLING BRAIN STRUCTURES FROM INDIVIDUAL CELL TYPES** |  |  |  |  |  |  |
| --- | --- | --- | --- | --- | --- | --- |
| **Study citation** | **Acronym** | **Tumor architecture and heterogeneity** | **Vascular cells** | **Immune cells** | **Oxygen level** | **Glucose concentration** |
| (Tang *et al.*, 2020) |  | **3D bioprinting** of a hyaluronic acid-rich hydrogel | n/a | THP-1 monocytes → M2 macrophages; primary human macrophages | Atmosphere | 16.7 mM (50% Neurobasal, 25% RPMI 1640, 25% Astrocyte Medium [ScienCell]) |
| (Seo *et al.*, 2022) |  | **Cells comprising the human blood-brain barrier** within a 3D hydrogel matrix BBB chip; GBM spheroids | Brain vascular pericytes; brain microvascular endothelial cells; astrocytes | n/a | Atmosphere | 5.6 mM (Endothelial Cell Medium, ScienCell) |
| (Straehla *et al.*, 2022), (Campisi *et al.*, 2018) |  | **Blood-brain barrier vascular network** within a microfluidic device, GBM spheroids | Human iPS-endothelial cells; pericytes; astrocytes | n/a | Atmosphere | 5.8 mM (VascuLife EnGS-Mv Medium, LifeLine Cell Technology) |
| (Stanton *et al.*, 2025) | **miBrain** | **Constructed with differentiated iPSCs for six major CNS cell types** assembled in 3D “Neuromatrix Hydrogel” and studied at 1-4 weeks in culture | Brain microvascular endothelial cells (BMECs); pericytes | Microglia-like cells | Atmosphere | 8.3 mM (Human Endothelial Serum Free Medium, Gibco) |

KEY RESOURCES TABLE.

KEY RESOURCES TABLE
| REAGENT or RESOURCE | SOURCE | IDENTIFIER |
| --- | --- | --- |
| Antibodies |  |  |
| Rabbit Monoclonal anti- EGFR (D38B1) | Cell Signaling | 4267L |
| Goat Polyclonal anti- IL13Rα2 | R&D Systems | AF146 |
| Mouse Monoclonal anti- ERBB 2 (3B5) | Thermo Fisher | MA5-13675 |
| Rabbit Polyclonal anti- von Willebrand Factor | Dako | A0082 |
| Mouse Monoclonal anti- CD31 (HEC7) | Invitrogen | MA3100 |
| Mouse Monoclonal anti- ZO- 1 (ZO1-1A12) | Invitrogen | 33-9100 |
| Rabbit Monoclonal anti- CD3 (SP162) | Abcam | ab135372 |
| Mouse Monoclonal anti- CD3 (OKT3) | Invitrogen | MA1-10175 |
| Rabbit Monoclonal anti- Cleaved Caspase 3 (5A1E) | Cell Signaling | 9664S |
| CODEX conjugated anti- EGFR (D38B1, BX021) (purchased from Cell Signaling and conjugated by Akoya Biosciences) | Cell Signaling, Akoya Biosciences | 26038SF |
| CODEX conjugated anti-IL13Rα2 (EPR22978-163, BX036) | Abcam, Akoya Biosciences | ab263874 |
| CODEX conjugated anti-HER2 (24D2, BX030) | Santa Cruz Biotechnology, Akoya Biosciences | sc-23864 |
| CODEX conjugated anti-SOX2 (RM427, BX027) | RevMAb, Akoya Biosciences | 31-1314-00 |
| CODEX conjugated anti-GFAP (SMI24, BX007) | BioLegend, Akoya Biosciences | 837404 |
| CODEX conjugated anti-BLBP<br>(EPR24033-13, BX042) | Abcam, Akoya<br>Biosciences | ab279652 |
| CODEX conjugated anti-DCX<br>(EPR19997, BX024) | Abcam, Akoya<br>Biosciences | ab222921 |
| CODEX conjugated anti-NESTIN<br>(EPR1301(2), BX004) | Abcam, Akoya<br>Biosciences | ab242400 |
| CODEX conjugated anti-S100β<br>(E7C3A, BX054) | Cell Signaling,<br>Akoya Biosciences | 42397 |
| CODEX conjugated anti-OLIG2<br>(EPR2673, BX035) | Abcam, Akoya<br>Biosciences | ab220796 |
| CODEX conjugated anti-COL1A1<br>(E3E1X, BX135) | Cell Signaling,<br>Akoya Biosciences | 66154 |
| CODEX conjugated anti-IBA1<br>(EPR16589, BX040) | Abcam, Akoya<br>Biosciences | ab178847 |
| CODEX conjugated anti-IBA1<br>(E83-77, BX019) | Abcam, Akoya<br>Biosciences | ab32042 |
| Anti-Hu CD31(AKYP0047)-BX001 | Akoya Biosciences | 4250019 |
| Anti-Hu SMA(AKYP0081)-BX013 | Akoya Biosciences | 4450049 |
| Anti-Hu CD3e(AKYP0062)-BX045 | Akoya Biosciences | 4550119 |
| Anti-Hu CD68(AKYP0050)-BX015 | Akoya Biosciences | 4550113 |
| Anti-Hu CD163(AKYP0114)-<br>BX069 | Akoya Biosciences | 4250079 |
| Anti-Hu CD14(AKYP0079)-BX037 | Akoya Biosciences | 4450047 |
| Anti-Hu/Mu Ki67(AKYP0052)-<br>BX047 | Akoya Biosciences | 4250019 |
| Anti-Hu CD20(AKYP0049)-BX064 | Akoya Biosciences | 4450094 |
| Alexa Fluor 488 Donkey Anti<br>Mouse IgG | Invitrogen | A21202 |
| Alexa Fluor 647 Donkey Anti<br>Mouse IgG | Invitrogen | A31571 |
| Alexa Fluor 555 Donkey Anti<br>Mouse IgG | Invitrogen | A31570 |
| Alexa Fluor 488 Goat Anti Mouse IgG2a | Invitrogen | A21131 |
| Alexa Fluor 647 Goat Anti Mouse IgG2B | Invitrogen | A21242 |
| Alexa Fluor 647 Goat Anti Mouse IgG1 | Invitrogen | A21240 |
| Alexa Fluor 555 Donkey Anti Rabbit IgG | Invitrogen | A31572 |
| Alexa Fluor 488 Donkey Anti Rabbit IgG | Invitrogen | A21206 |
| Alexa Fluor 555 Goat Anti Rabbit IgG | Invitrogen | A21429 |
| Alexa Fluor 647 Donkey Anti Goat IgG | Invitrogen | A21447 |
| Alexa Fluor 488 Streptavidin | Invitrogen | S32354 |
| Alexa Fluor 647 Streptavidin | Invitrogen | S32357 |
| Alexa Fluor™ 750-RX007 for PhenoCycler (CODEX) | Akoya Biosciences | 6450003 |
| Alexa Fluor™ 647-RX030 for PhenoCycler | Akoya Biosciences | 6550020 |
| Alexa Fluor™ 750-RX001 for PhenoCycler | Akoya Biosciences | 6450001 |
| Atto 550-RX035 for PhenoCycler | Akoya Biosciences | 6250010 |
| Alexa Fluor™ 647-RX054 for PhenoCycler | Akoya Biosciences | 6550029 |
| Alexa Fluor™ 750-RX064 for PhenoCycler | Akoya Biosciences | 6450028 |
| Atto 550-RX135 for PhenoCycler | Akoya Biosciences | 6250050 |
| Alexa Fluor™ 647-RX045 for PhenoCycler | Akoya Biosciences | 6550024 |
| Alexa Fluor™ 750-RX004 for PhenoCycler | Akoya Biosciences | 6450002 |
| Atto 550-RX069 for PhenoCycler | Akoya Biosciences | 6250027 |
| Alexa Fluor™ 647-RX021 for PhenoCycler | Akoya Biosciences | 6550028 |
| Alexa Fluor™ 750-RX019 for PhenoCycler | Akoya Biosciences | 6450007 |
| Atto 550-RX040 for PhenoCycler | Akoya Biosciences | 6250021 |
| Alexa Fluor™ 647-RX036 for PhenoCycler | Akoya Biosciences | 6550022 |
| Alexa Fluor™ 750-RX013 for PhenoCycler | Akoya Biosciences | 6450005 |
| Atto 550-RX037 for PhenoCycler | Akoya Biosciences | 6250017 |
| Alexa Fluor™ 647-RX042 for PhenoCycler | Akoya Biosciences | 5550015 |
| Atto 550-RX047 for PhenoCycler | Akoya Biosciences | 6250012 |
| Alexa Fluor™ 647-RX015 for PhenoCycler | Akoya Biosciences | 6550027 |
| Alexa Fluor™ 647-RX027 for PhenoCycler | Akoya Biosciences | 6550019 |
| Alexa Fluor™ 647-RX024 for PhenoCycler | Akoya Biosciences | 6550018 |
| PE-Cy™7 Mouse anti- Human CD19 (SJ25C1) | BD Pharmingen™ | 557835 |
| Bacterial and virus strains |  |  |
| Cltx-IgG4(L235E, N297Q)-CD28gg-Zeta(CO)-T2A - CD19t_epHIV7 (Clinical) | Dr. Christine Brown laboratory at City of Hope | vF1267 |
| IL13op-IgG4op(L235E, N297Q)-CD4tmop-41BB-Zetaop-T2A-CD19t_epHIV7(clinical) | Dr. Christine Brown laboratory at City of Hope | vF0455 |
| Biological samples |  |  |
| Human glioblastoma tissue | City of Hope Research Pathology Core | N/A |
| Human blood samples | City of Hope<br>Research Pathology<br>Core | N/A |
| Chemicals, peptides, and recombinant proteins |  |  |
| Chlorotoxin | Alomone Labs | RTC-450 |
| CLTX:biotin | Purchased from<br>Almone and<br>conjugated by the<br>Synthetic<br>Biopolymer<br>Chemistry Core at<br>City of Hope | N/A |
| Hoechst 33342 | Life Technologies | H3570 |
| Prolong™ Glass Antifade<br>Mountant | Invitrogen | P36984 |
| 10X Buffer for PhenoCycler | Akoya Biosciences | 7000001 |
| Assay Reagent for PhenoCycler | Akoya Biosciences | 7000002 |
| Nuclear Stain for PhenoCycler | Akoya Biosciences | 7000003 |
| Xylenes (Histological) | Fisher Scientific | X3P-1GAL |
| Ethyl Alcohol 100% (200 proof) | Pharmco by<br>Greenfield Global | 111000200 |
| 10X Citrate Buffer, pH=6.0 0.1M | Sigma-Aldrich | C9999-100ML |
| 16% Paraformaldehyde | Electron Microscope<br>Sciences | 15710 |
| PBS 1X (Dulbecco's Phosphate<br>Buffered Saline Solution) | FUJIFILM Irvine<br>Scientific | 9236 |
| Poly-L-lysine 0.1% | Sigma- Aldrich | P8920 |
| Nuclease-Free Water | Thermo Fisher<br>Scientific | AM9938 |
| Methanol | Sigma- Aldrich | 34860-1L-R |
| BrainPhys™ Neuronal Media | Stem Cell<br>Technologies | 05790 |
| GlutaMAX™ Supplement (100X) | Gibco | 35050061 |
| MEM Non-Essential Amino Acids Solution (100X) | Gibco | 11140050 |
| N-2 Supplement (100X) | Gibco | 17502048 |
| B-27™ Supplement (50X), minus vitamin A | Gibco | 12587010 |
| 2-Mercaptoethanol (1000X) | Gibco | 21985023 |
| Insulin solution human (9.5-11.5 mg/ml) | Sigma-Aldrich | I9278 |
| Matrigel® Growth Factor Reduced (GFR) Basement Membrane Matrix, Phenol Red-free, LDEV-free | Corning | 356231 |
| Penicillin-Streptomycin (10,000 U/mL) (100X) | Gibco | 15140122 |
| NuSieve™ 3:1 Agarose | Lonza | 50090 |
| TopVision Low Melting Point Agarose | Thermo Scientific | R0801 |
| Tissue- Tek® O.C.T. Compound | Sakura | 4583 |
| X- VIVO™ 15 | Lonza | BEBP04-744Q |
| Fetal Bovine Serum (Defined, FBS) | Cytiva | SH30070.03 |
| Transnet YX Tissue Hibernat® A | BrainBits | HA |
| Amphotericin B 250 µg/ml | Gibco | 15290-026 |
| 10X TBS | Bioland Scientific LLC | TBS01-02 |
| CAS-Block™ | Invitrogen | 008120 |
| Paraformaldehyde | Sigma- Aldrich | P6148-500G |
| PBS pH7.4 (10X) | Gibco | 70011-044 |
| Fluorescein diacetate | Sigma-Aldrich | F7378-5G |
| Propidium iodide | Tocris Bioscience | 5315 |
| Sodium Azide (NaN<sub>3</sub>) | Sigma- Aldrich | S8032 |
| HBSS (1X) | Gibco | 14175-095 |
| UltraComp eBeads™ Plus Compensation Beads | Invitrogen | 01-3333-42 |
| Dynabeads® Human T- Expander | Invitrogen | 11141D |
| ViaStain™ AOPI staining solution | Nexcelom Bioscience | CS2-0106-25ml |
| DMEM (1X, Dulbecco's Modified Eagle Medium) | Gibco | 11960-051 |
| PEIpro® | Polyplus | 101000017 |
| Sodium butyrate ≥98.5% (GC) | Gibco | B5887-5g |
| D-(+)-Lactose monohydrate | Avantor by VWR | JT2249-1 |
| RPMI 1640, 1X | Corning | 10-041-CM |
| Human Serum Albumin (HSA), 25% Flexbumin (50 mL, bag) | City of Hope Pharmacy | NDC 0944-0493-01 |
| 50U/ml IL2 | CellGenix | 1013-050 |
| 0.5ng/ml IL15 | City of Hope Pharmacy | NDC 76310-0022-01 |
| Protamine Sulfate | City of Hope Pharmacy | NDC 63323-0229-15 |
| DAPI, FluoroPure™ grade | Invitrogen | D21490 |
| Hexadimethrine bromide (polybrene) | Sigma- Aldrich | H9268-10G |
| Buffer RLT Plus RNeasy® Plus lysis buffer | Qiagen | 1048449 |
| Critical commercial assays |  |  |
| EZ-Link™ Sulfo-NHS-LC-Biotinylation Kit | Thermo Scientific | 21435 |
| Staining Kit for PhenoCycler | Akoya Biosciences | 7000008 |
| RNeasy® Mini Kit (50) | Qiagen | 74104 |
| Qubit RNA HS Assay Kit | Fisher Scientific | Q32852 |
| Kapa RNA HyperPrep Kit with RiboErase | Roche | KK8560 |
| Deposited data |  |  |
| Bulk RNA sequencing | This paper |  |
| Experimental models: Cell lines |  |  |
| HEK 293T cells | City of Hope, Center for Biomedicine and Genetics | cJ03756 |
| Jurkat Cells | ATCC | TIB-152 |
| Human Tn/mem T cells | Dr. Christine Brown laboratory at City of Hope | N/A |
| Experimental models: Organisms/strains |  |  |
| Organotypic slice culture | This paper |  |
| Recombinant DNA |  |  |
| pCHGP-2 | Dr. Jiing-Kuan Yee laboratory at City of Hope | pJ01506 |
| pCMV-Rev2 | Dr. Jiing-Kuan Yee laboratory at City of Hope | pJ01505 |
| pCMV-G | Dr. Jiing-Kuan Yee laboratory at City of Hope | pJ01507 |
| Cltx-IgG4(L235E, N297Q)-CD28gg-Zeta(CO)-T2A - CD19t_epHIV7 (Clinical) | Dr. Christine Brown laboratory at City of Hope | pF02932 |
| IL13op-IgG4op(L235E, N297Q)-CD4tmop-41BB-Zetaop-T2A-CD19t_epHIV7(clinical) | Dr. Christine Brown laboratory at City of Hope | pF02630 |
| Software and algorithms |  |  |
| Imaris | Bitplane | https://imaris.oxinst.com |
| ImageJ 1.53c | NIH | https://imagej.nih.gov/ij |
| QuPath 0.4.4 | QuPath developers,<br>The University of<br>Edinburgh | https://qupath.github.io/ |
| GraphPad Prism | GraphPad Software | https://www.graphpad.com |
| Adobe Illustrator 2025 | Adobe | https://www.adobe.com/products/illustrator |
| FCS Express | De Novo Software<br>by Dotmatics | https://denovosoftware.com/ |
| Zeiss Zen 3.11 | Carl Zeiss | https://www.zeiss.com/microscopy/us/products/software/zeiss-zen.html |
| NDP. view 2.7.25 | Hamamatsu<br>Photonics K.K | U12388-01 |
| GBMDeconvoluter | Ajaib et.al 2022;<br>Ruiz-Moreno et.al<br>2022 | https://gbmdeconvoluter.leeds.ac.uk/ |
| Cellometer K2 | Nexcelom<br>Bioscience LLC |  |
| R Project | Open Source | https://www.r-project.org/ |
| RStudio | Posit | https://posit.co/download/rstudio-desktop/ |
| trimmomatic v.0.39 | Illumina | https://github.com/usadellab/Trimmomatic |
| fastp v.0.23.2 | OpenGene | https://github.com/OpenGene/fastp |
| STAR v.2.7.9a | Dobin et al., 2012 | https://github.com/alexdobin/STAR |
| Subread v.2.0.3 | Liao et al., 2013 | https://subread.sourceforge.net/ |
| edgeR v.3.32.1 | Chen et., al 2025 | https://bioconductor.org/packages/release/bioc/html/edgeR.html |
| org.Hs.eg.db v.3.17.0 | Marc Carlson,<br>Bioconductor open<br>source | DOI: 10.18129/B9.bioc.org.Hs.eg.db |
| sva v. 3.48.0 | Jeffrey T. Leek et<br>al., Bioconductor<br>open source | DOI: 10.18129/B9.bioc.sva |
| Gene Set Enrichment Analysis v4.3.3 | Subramanian, Tamayo, et al. (2005, PNAS); Mootha, Lindgren, et al. (2003, Nature Genetics). | https://www.gsea-msigdb.org/gsea/index.jsp |
| Other |  |  |
| Glass Cover Slips #1<sup>1/2</sup> 22x22mm | Electron Microscopy Sciences | 72204-01 |
| Superfrost Plus Microscope Slides | Fisher Scientific | 12-550-15 |
| Cell Culture Inserts 0.4µm, 30mm Diameter | Millicell® | PICM0RG50 |
| Leica VT1000 S Vibrating blade microtome | Leica Biosystems | N/A |
| Tissue- Tek® Cryo3 Flex ® Model 6200 | Sakura | N/A |
| Nalgene™ Rapid- Flow™ 75mm Filter Unit | Thermo Scientific | 156-4020 |
| Globe Scientific 4oz. Specimen Containers | Fisher Scientific | 22-170-143 |
| Corning® 150ML Filter System | Corning | 431155 |
| DynaMag-50 magnet | Invitrogen | 12302D |
| Krazy Super Glue | Krazy Glue | N/A |
| Sapphire Blade, 1.5" (40mm) | Ted Pella, Inc. | 121-18 |
| Nanodrop One | Fisher Scientific | ND-ONE-W |
| Cellometer K2 | Nexcelom Bioscience | CMT-K2-MX-150 |
| NovaSeq 6000 | Illumina |  |
| TintoRetriever pressure cooker | BIO SB | NC1709641 |
| Cellometer cell counting slides | Revvity | SD100 |

